# Islet Gene View - a tool to facilitate islet research

**DOI:** 10.1101/435743

**Authors:** Olof Asplund, Petter Storm, Vikash Chandra, Emilia Ottosson-Laakso, Gad Hatem, Dina Mansour-Aly, Ulrika Krus, Hazem Ibrahim, Emma Ahlqvist, Tiinamaija Tuomi, Erik Renström, Olle Korsgren, Nils Wierup, Claes Wollheim, Isabella Artner, Hindrik Mulder, Ola Hansson, Timo Otonkoski, Leif Groop, Rashmi B Prasad, on behalf of the Human Tissue Laboratory at Lund University Diabetes Centre

## Abstract

Changes in the hormone-producing pancreatic islets are central culprits in type 2 diabetes (T2D) pathogenesis. Characterization of gene expression in islets how it is altered in T2D are therefore vital in understanding islet function and T2D pathogenesis. We leveraged RNA-sequencing and genome-wide genotyping in islets from 188 donors to create the Islet Gene View (IGW) platform to make this information easily accessible to the scientific community. The IGW combines expression data for a given gene with phenotypical data such as T2D status, BMI, HbA1c, insulin secretion, purity of islets, etc.), regulation of gene expression by genetic variants e.g., expression quantitative trait loci (eQTLs) and relationship with expression of islet hormones. In IGW, 285 differentially expressed genes (DEGs) were identified in T2D donors islets compared to controls. Forty percent of the DEGs showed cell-type enrichment and a large proportion of them were significantly co-expressed with islet hormone-encoding genes like glucagon (*GCG*, 56%), amylin (*IAPP*, 52%), insulin (*INS*, 44%) and somatostatin (*SST*, 24%). Inhibition of two DEGs, *UNC5D* and *SERPINE2* impaired glucose-stimulated insulin secretion and impacted cell survival in a human beta-cell model.

**Significance Statement:** We present Islet Gene View (IGW), a web resource facilitating information on gene expression in human pancreatic islets from organ donors easily accessible to the scientific community. In IGW, we explored RNA expression from 188 donor-islets and examined their relationship with islet phenotypes including insulin secretion and expression of genes encoding islet hormones. GWAS have shown 403 genetic variants associated with risk of type 2 diabetes (T2D) risk, however, the target genes and function of these variants in islets are largely unknown. By linking T2D risk variants to expression in islets from T2D and non-diabetic donors as well as islet phenotypes, use of IGW provided new insight into mechanisms by which variants in these loci may increase risk of T2D.

## Introduction

Type 2 diabetes (T2D) is a chronic condition that arises from the inability of the body to maintain glucose homeostasis. It has broadly been attributed to inability of insulin secretion to compensate for insulin resistance, the latter often arising due to obesity and physical inactivity (1). Genome-wide association studies (GWAS) have identified 403 loci robustly associated with predisposition to T2D, or related glycemic traits (>70) (1–3). A vast majority of the loci primarily influence insulin secretion (3, 4), thus emphasizing the central role of the islets of Langerhans and beta-cell dysfunction in T2D pathogenesis.

Gene expression analysis provides a possibility to link between genetics and cellular function and is crucial for the elucidation of pathophysiological mechanisms. Information on gene expression in different tissues has greatly served the understanding of disease mechanisms. For example, the Genotype-Tissue Expression (GTEx) project (5) is a pioneering example on how to share such information from deceased humans. Unfortunately, GTEx has limited information on human pancreatic islets of Langerhans, as RNA sequencing was performed on whole pancreas, not islets, and only from a limited number of deceased donors. Importantly, the pancreas needs to be removed while blood flow is still intact to retain functionality of the pancreatic cells (6), and therefore more information on human pancreatic islets can be derived from organ donors selected for transplantation purposes where blood flow has been kept intact until pancreas excision and islet isolation is possible.

One of the main objectives of the EU-funded strategic research area, Excellence of Diabetes Research in Sweden (EXODIAB), is to facilitate diabetes research globally, e.g., by creating resources and tools that can be used by the research community. One of its central platforms, The Human Tissue Laboratory (HTL), has generated a large repository of tissue samples (human pancreatic islets, fat, liver and muscle) from deceased organ donors. It comprises gene expression (bulk RNA sequencing) and genome-wide genetic variation data (GWAS) from human pancreatic islets, as well as some of the target tissues for insulin. Here we describe the development of a web-based tool, the Islet Gene View (IGW), which will allow rapid and robust overview of data, as well as “look-up” of genes of interest. The underlying database shows differences in gene expression between T2D and non-diabetic donors and their relationship with specific islet phenotypes as well as the effects of genetic variation on gene expression, *i.e.* eQTLs (Expression Quantitative Trait Loci). The tool includes some well validated examples to reinforce the usefulness and precision of the tool.

## Results

### Islet Gene View (IGW) – a web resource for gene expression in human pancreatic islets

We created the Islet Gene View (IGW) web tool to functionally annotate all genes expressed in human pancreatic islets and to provide a platform to look up genes of interest. Islet Gene View (IGW) is accessible at https://mae.crc.med.lu.se/IsletGeneView/ after registration. IGW uses several common gene identifiers (*e.g.* gene symbol, Ensembl gene ID, and full gene name), and provides graphs of gene expression in relation to islet phenotypes and expression of other genes (Supplementary figure 1). An example graph is given in figure 1. The first graph reveals gene expression in human pancreatic islets and other target tissues for insulin (e.g., a 12-donor tissue panel of biopsies from fat, liver and skeletal muscle). The second graph shows the relationship between gene expression and purity (islet volume fraction, i.e the proportion of endocrine component over exocrine), followed by its expression in relation to other genes expressed in the islets. Subsequent figures show gene expression in relation to glycemic islet phenotypes such T2D, glucose tolerance defined by HbA1c strata (normal glucose tolerance, NGT: HbA1c <6%, impaired glucose tolerance, IGT: HbA1c between 6-6.5%, and T2D: HbA1c > 6.5%), HbA1c as a linear variable as well as BMI, and glucose-stimulated insulin secretion (GSIS) in islets (stimulatory index, SI). Of interest is also co-expression with other islet cell-specific genes such as *INS, GCG* and *IAPP*. The Ensembl gene ID is the primary identifier for the expression database. Additional gene identifiers and other annotations were derived from the gencode v22 gene transfer format (GTF) annotation file.

**Figure 1:**
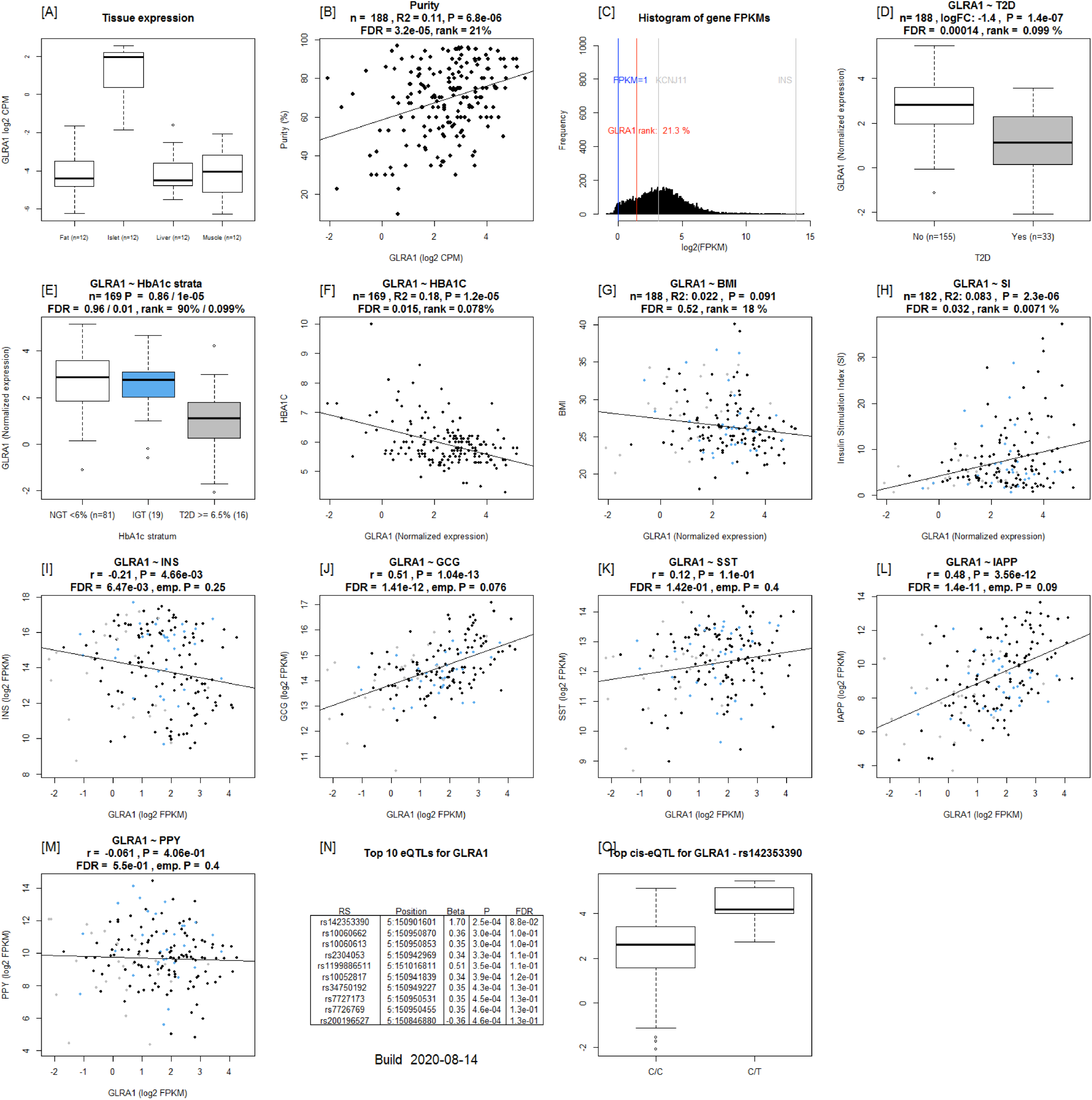
Example output from Islet Gene View of *GLRA1* [A]: expression of the gene in fat, islets, liver, and muscle in the same pool of 12 individuals. [B]: Gene expression as a function of purity, defined as the percentage of endocrine tissue. [C]: Expression of the selected gene in relation to other genes in islets. [D-H]: Gene expression in relation to several diabetes-related phenotypes, i.e. T2D diagnosis (figure 1D), HbA1c stratum (1E), continuous HbA1c (1F), BMI (1G), and stimulatory index (1H). Test statistics are reported, namely: coefficient of determination (R2), nominal P-value, and percentage rank among all genes as calculated based on sorted P-values. (46): Gene expression in relation to the secretory genes INS [I] GCG [J], SST [K], PPY [L], and IAPP [M]. Spearman’s ρ (r) and the P-value of the gene based on the empirical correlation distribution is reported. INS = Insulin, GCG = glucagon, SST = somatostatin, PPY = pancreatic polypeptide, IAPP = islet amyloid polypeptide.

### Multiple genes are differentially expressed between islets from type 2 diabetic and nondiabetic donors

IGW, in the current version, is based on global gene expression in human pancreatic islets obtained from 188 organ donors (Supplementary Table 1). In the browser, a differential expression analysis of 33 donors with a clinical diagnosis of T2D and 155 diabetes-free individuals (Supplementary Figure 1) showed that expression of 285 out of a total of 14 108 genes differed significantly between the two groups (FDR <= 0.05, Figure 2i, Supplementary Table 2). Of these differentially expressed genes (DEGs), expression of 120 genes was downregulated whereas that of 165 was upregulated in T2D donor islets (Figure 2i). Expression of 24 genes showed significant correlation with HbA1c levels. The expression levels of genes that were upregulated in T2D donors were also positively correlated with HbA1c whereas those of downregulated genes were negatively correlated (Supplementary Table 3, Figure 1).

**Figure 2.**
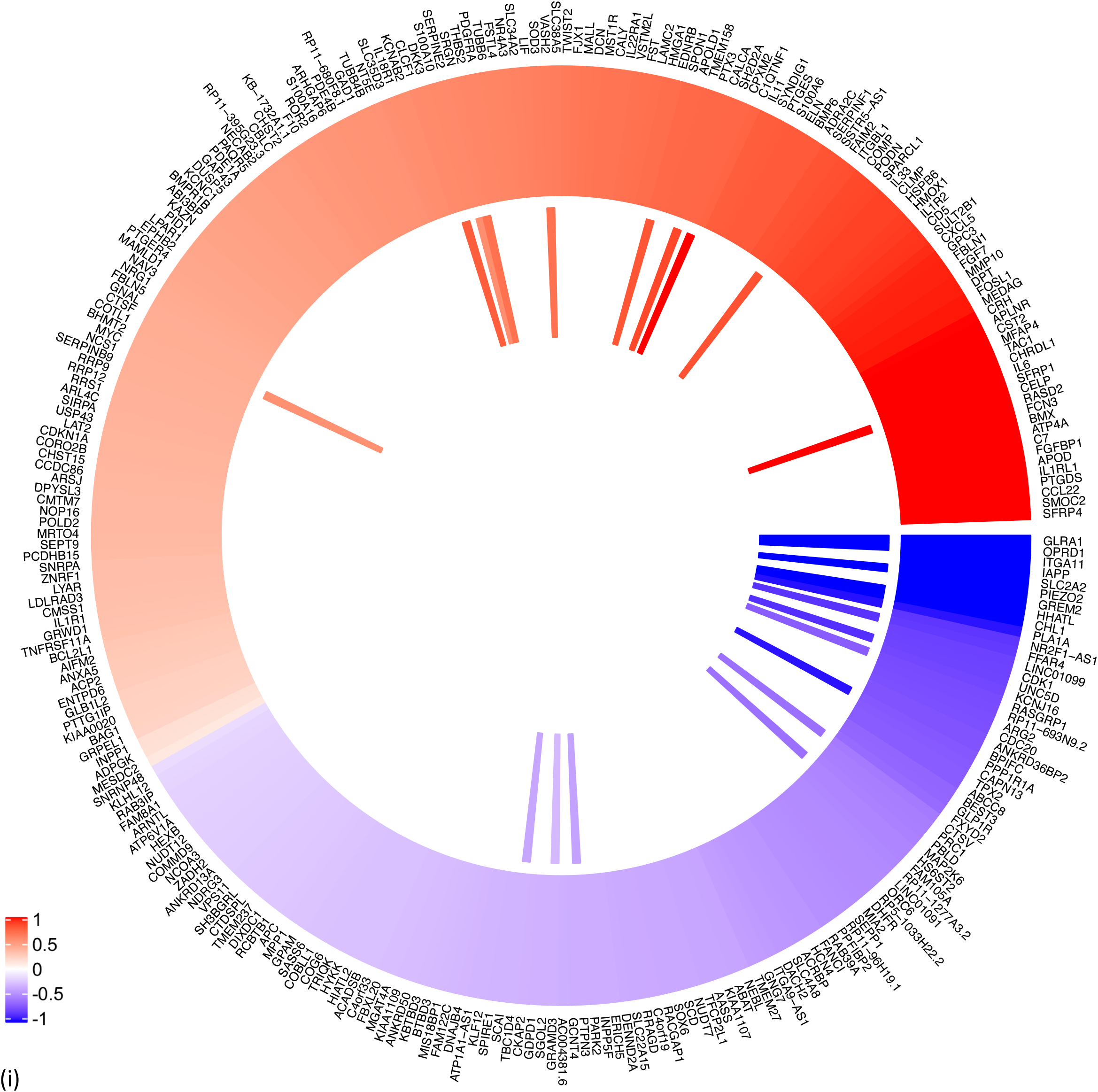
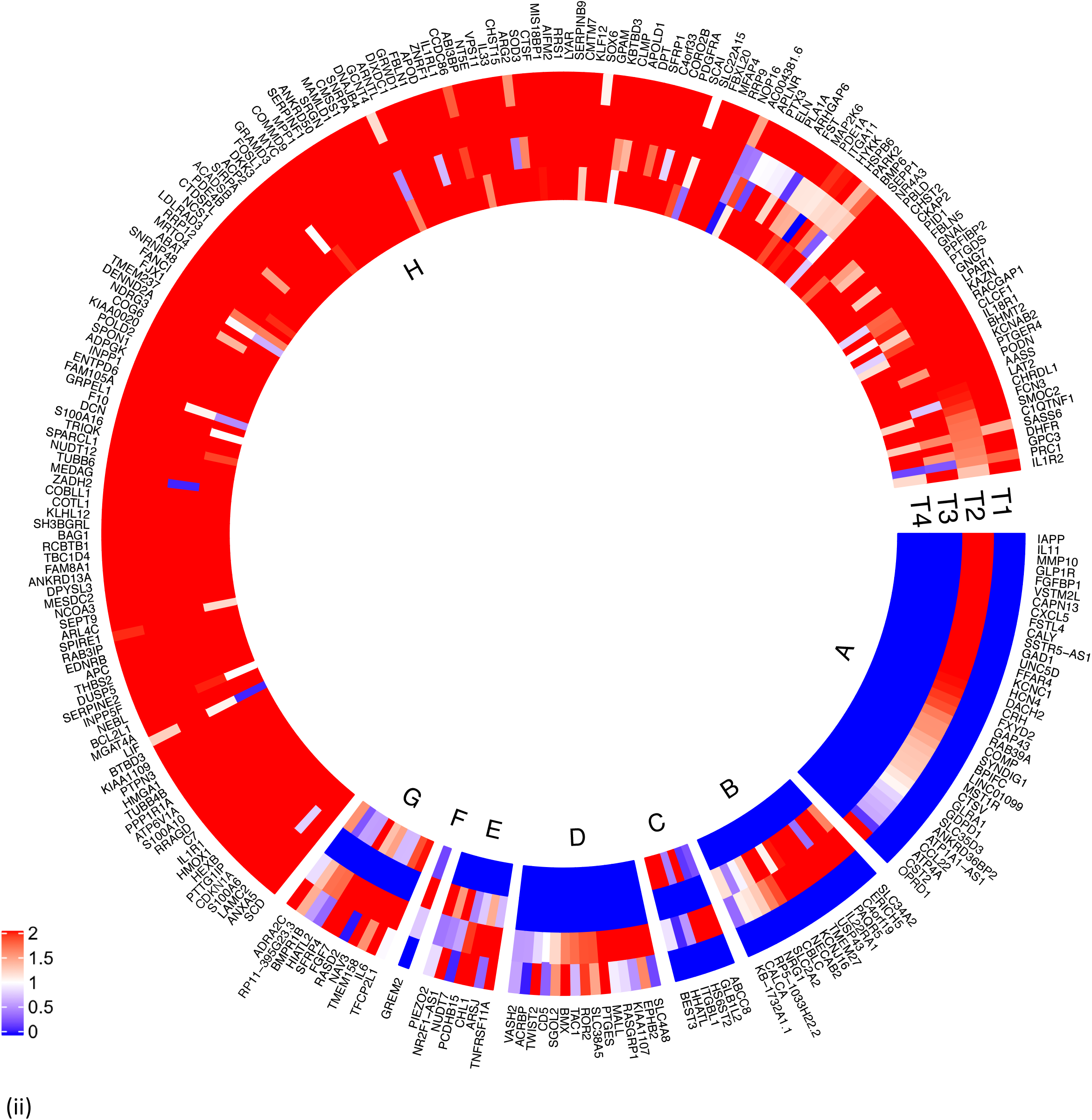
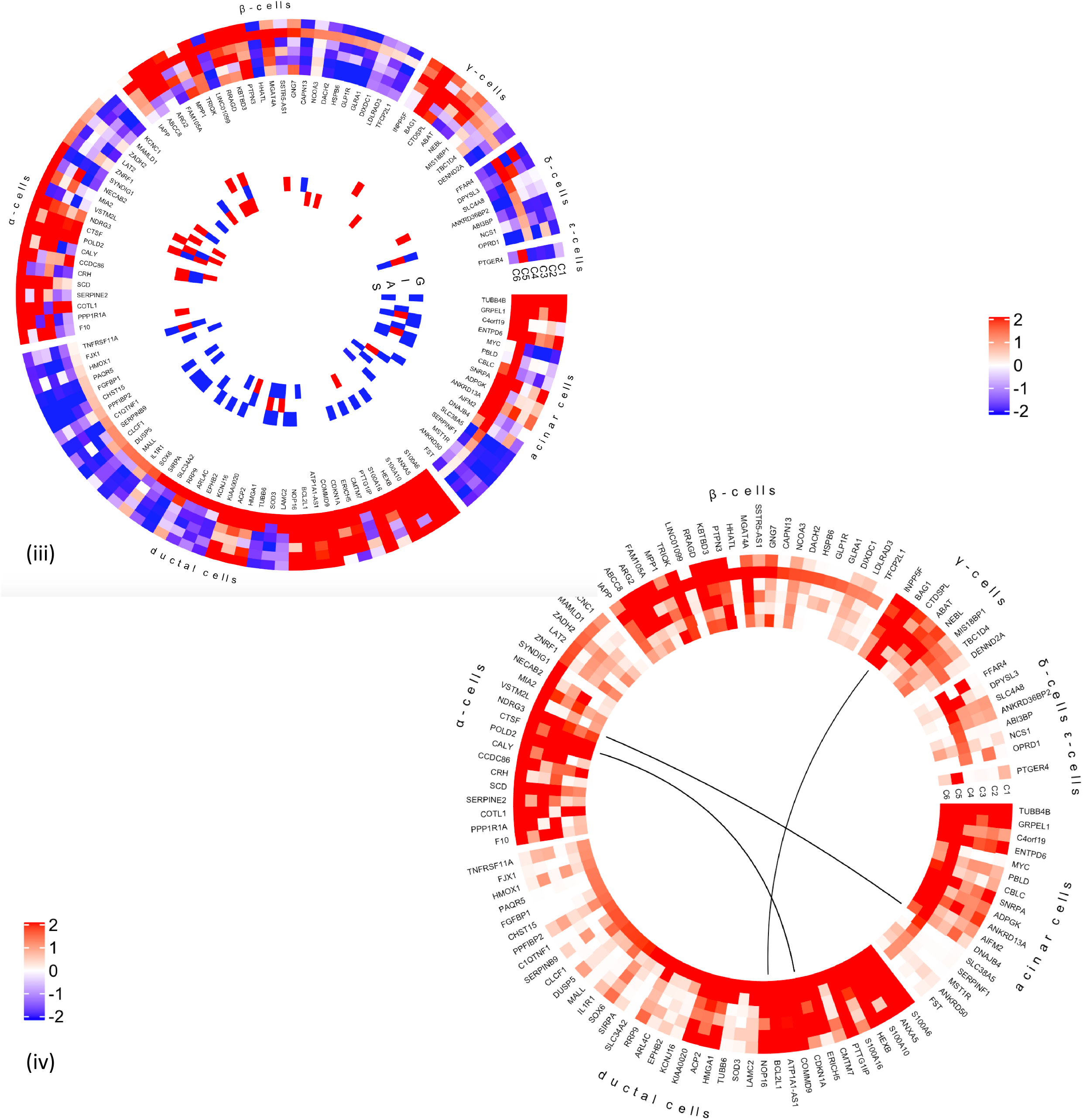
Characterization of differentially expressed genes (DEGs) between islets from T2D donors compared to controls. (i) Outer track: 120 genes were down regulated (indicated in blue) whereas 165 genes were upregulated in islets from T2D donors (indicated in red). Inner track: Genes showing significant positive correlation with HbA1c levels (red) and negative correlation (blue). (ii) Expression of the DEGs in fat (F), islet (I), liver (L) and muscle (M). T1 shows expression of DEGs in Fat, T2 in Islets, T3 in liver and T4 in muscle. Expression was defined as ≥ 1 count per million (CPM). DEGs expressed in islets not in other tissues) are shown in Segment A, islet and liver in Segment B, islet and muscle in Segment C, fat and islet in Segment D, fat, islet and liver in segment E, islet, liver and muscle in segment F, fat islet and muscle in segment G, and all 4 tissues in segment H. Majority of the DE genes were expressed in all 4 tissues. coded as blue < 1, 0 = white, red ≥ 1. (iii) DEGs, cell type enrichment and correlation with secretory genes. DEGs which are enriched in specific islet cell types are separated into specific segments Outer tracks show expression in scRNAseq ^1^. Outermost track show expression in alpha cells (C1), followed by beta cells(C2), gamma cells (C3), delta cells (C4), acinar cells (C5) and ductal cells (C6). Mean of RPKM (log2) values are plotted, with values code as: blue < 1< red. Inner tracks show correlation with secretory genes starting with GCG on the outside (G), followed by INS (I), IAPP (A) and SST (S), coded as - 0.5 >= blue, 0 = white, 0.5 <= red. (iv) DEGs grouped by cell type enrichment as reported in Segerstolpe et al. The links show networks as inferred by GeNets ^2^.

To validate the data that underlie the IGW, we sought to replicate our key findings by comparing our list of DEGs with previously published data. For this, we selected a study presenting DEG data from microarray experiments (7), one of which was run on islets obtained from partially pancreatectomized patients and a second set of experiments carried out on islets from organ donors. Nine genes were replicated with directional consistency with our dataset in both the microarray sets, including *UNC5D, PPP1R1A, TMEM37, SLC2A2, ARG2, CAPN13, FFAR4, HHATL* and *CHL1*. Expression changes in 13 genes were replicated in the expression data obtained from organ donors and that of 15 genes in the dataset of the partially pancreatectomized donors (Figure 3). Of the total 39 replicated genes, 27 also associated with HbA1c levels (Supplementary table 3). Of particular interest were the genes *HHATL, CHL1* and *SLC2A2*, the expression of which was concordant to findings from a previous study including a subset of the current islets (8) (Supplementary Table 2). The expression of all the above 3 genes correlated nominally with stimulatory index (measure of GSIS in the islets) as well as with BMI. The expression of *HHATL* correlated with expression of *INS, SST* and *IAPP, CHL1* with *GCG* and *SLC2A2* with *IAPP* (Supplementary figure 2). We and others have previously shown that a genetic variant at the *CHL1* locus (rs9841287 SNP) is associated with fasting insulin concentrations (9, 10). In addition, this locus was an eQTL for the *CHL1* and *CHL1-AS1* genes (beta = 0.29 and 0.27, P = 0.028 and 0.04 respectively, increasing allele = G).

**Figure 3.**
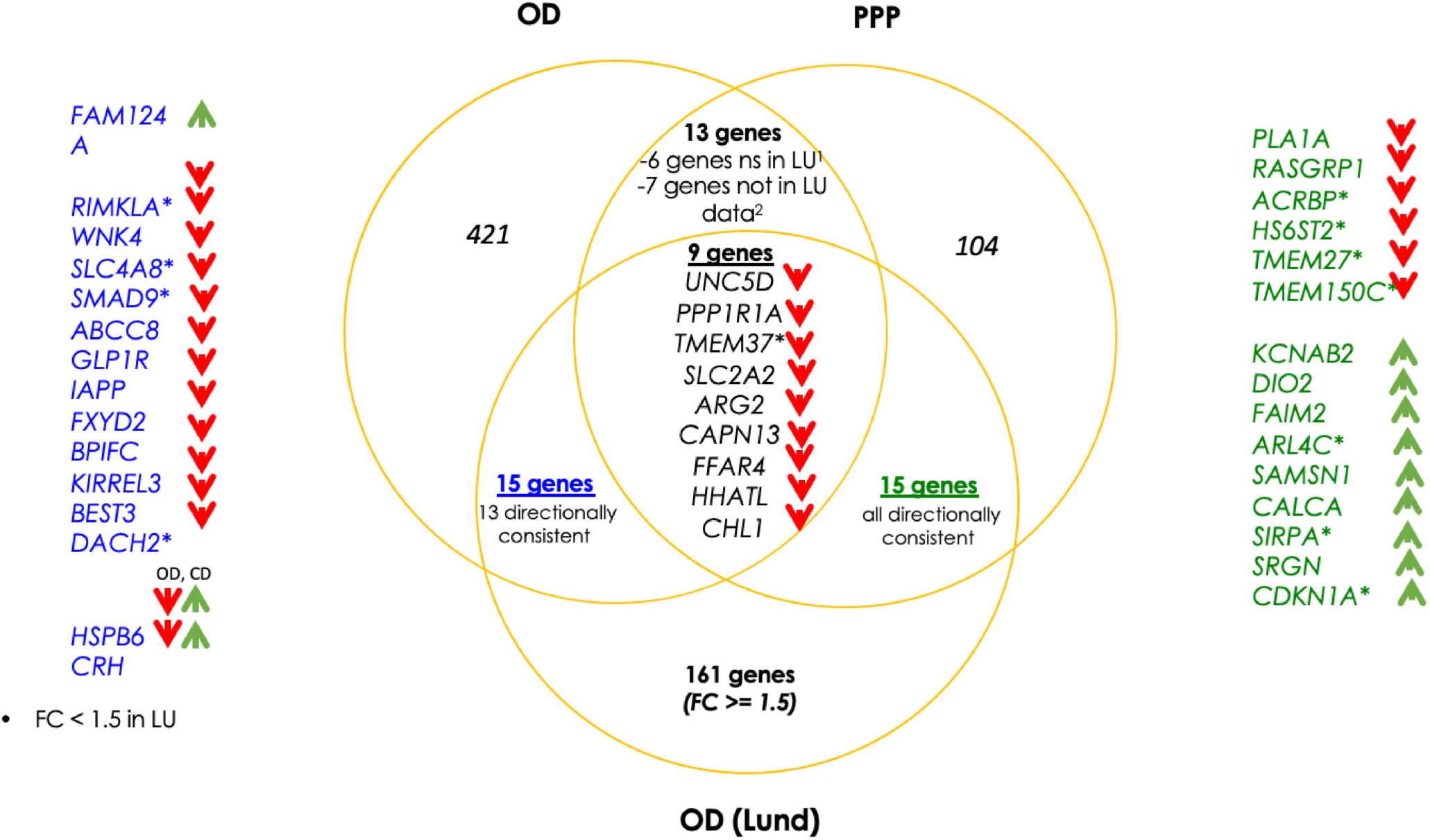
Replication of differentially expressed genes (DEGs) in comparison with data from Solimena et al^3^. Venn diagram shows the overlap of DEGs from each of the studies. OD and PPP show DE data from organ donor islets and partially pancreatectomized donor islets from Solimena et al whereas OD (LUND) show DE genes from our data. 9 genes were replicated in all 3 data sets. 13 genes were replicated between the OD islets from Solimena et al (presented in blue on the left) and our data whereas 15 genes were replicated between PPP islets and our data (presented in green on the right). The arrows indicate the direction of effect: red arrows pointing down show down regulation in T2D islets whereas green arrows pointing up are upregulated.

Of the DEGs, the expression of *GLRA1*, was significantly associated with the stimulatory index (Figure 1). Another gene of interest was *FXYD2*, the expression of which was downregulated in T2D donor islets and correlated with HbA1c levels. Its expression was also downregulated in beta-cells from T2D donors in a previous study using single cell RNAseq (11).

### Changes in gene expression which are potentially the cause or consequence of hyperglycemia

Dysregulated gene expression in islets from T2D donors can be causal in T2D pathogenesis or be secondary to the long-term hyperglycemia. To identify changes in genes which are causal and those which develop as a consequence of hyperglycemia, we next compared our set of differentially expressed genes with our previously published study (12) in which we treated normo- (n = 31) and hyperglycemic (n = 14) islets with high (18.9mmmol/L) and low (5.5mmol/L) glucose respectively. The expression of 13 genes was unaffected by short-term glucose exposure (acute hyperglycemia-HG), but altered due to long-term hyperglycemia (12). The expression of these genes also correlated with insulin secretion with consistent directionality in the current study(i.e. increased expression in HG, negative correlation with insulin secretion and vice versa) (Supplementary Table 4). We hypothesize that changes in expression of these genes are likely causal for T2D pathogenesis.

#### A majority of the genes expressed in islets and DEGs show variable expression in fat, liver and muscle

Gene expression is a means by which a genome controls cell differentiation and consequently development of different tissues. Gene expression and its genetic regulation have been determined to vary in its specificity; it ranges from being highly tissue specific to ubiquitous, the latter facilitating a higher degree of pleiotropy. The extent to which islet-expressed genes found in IGW also show expression in other tissues, especially those of relevance to T2D is yet to be undetermined. To address this, we examined whether genes expressed in islets were also expressed in fat, liver and muscle obtained from the same individuals (defined as ≥ 1cpm in >80% of samples). Indeed, we found a large proportion of genes (11118) to be expressed in all four tissues at varying levels (Supplementary figure 3). In addition, each tissue also had unique expression patterns: 1122 genes were expressed in islets, 713 in muscle, 529 in liver and 1112 in adipose tissue. Of the 285 DEGs, 36 genes showed highest expression in islets compared to the other tissues; this included the islet hormone coding *IAPP*, antisense of *SSTR5* coding *SSTR5-AS1, GLRA1, FXYD2* and *UNC5D*. Fifteen genes were expressed in islet and liver including *HHATL*, whereas 40 others were expressed in at least one other tissue (Figure 2ii). A majority of the genes seemed to be ubiquitously expressed in all tissues tested (Figure 2ii).

#### DEGs in T2D donor islets are significantly co-expressed with islet hormone encoding genes

Pancreatic islets comprise insulin-secreting beta-cells, alpha-cells which secrete glucagon, delta-cells which secrete somatostatin, F-cells (also called PP cells) which secrete pancreatic polypeptide Y and ghrelin cells which produce ghrelin. In addition, beta-cells also secrete islet amyloid polypeptide (*IAPP*), which was a DEG between T2D and controls. Therefore, genes found in IGW whose expression correlates with insulin and glucagon expression are likely to influence insulin expression or are potential downstream targets of *INS* and *GCG*. Altogether, 11238 genes were co-expressed (FDR < 0.05) with *INS* and 4519 of them were empirically significant (i.e. when gene-gene correlations for all genes expressed in islets were considered). The corresponding numbers for *GCG* were 9347 and 789, for *SST* were 9937 and 1172, whereas 10638 and 2610 for *IAPP*. The number of genes showing positive and negative co-expression with *INS, GCG, IAPP* and *SST* and overlapping correlations regardless of direction are shown in Supplementary figure 4 A-C respectively. A list of top 10 genes showing the strongest correlation with islet hormone encoding genes is provided in Supplementary table 5.

The expression of 124 DEGs correlated significantly with *INS* expression of which 24 were significant at the empirical level (Supplementary Table 6, Fig 1c). The corresponding numbers were 160 and 42 for *GCG* (Supplementary Table 7), 147 and 65 for *IAPP* (Supplementary Table 8), 67 and 8 for *SST* (Supplementary Table 9), none for *PPY* (Supplementary Table 10), and 0 and 2 for *GHRL* (Supplementary Table 11) (Figure 2iii).

#### Genes whose expression is altered in T2D donor islets show cellular heterogeneity in enrichment of expression

We next compared the DEG gene-set derived from IGW to a gene list of cell-type enriched genes from a previously published study applying single cell RNA sequencing (scRNAseq) of islets (11). This yielded an overlap of 20 genes enriched in alpha cells, 23 in beta-cells, 8 in gamma, 7 in delta, 17 in acinar and 39 in ductal cells. Of genes showing correlation with at least one islet hormone encoding gene, the largest proportion of genes was co-expressed with *GCG* (65%), followed by *IAPP* (57%), *INS* (50%) and *SST* (35%) (Fig 2iii). Two genes were co-expressed with *GHRL* (*PPP1R1A* and *FAM105A*) whereas there were no significant co-expression between the DEGs and *PPY*. The expression of 36 genes showed opposite correlation with *INS* and *GCG* (i.e. positive with *INS* and negative with *GCG*, and vice versa). These included the (i) *IAPP* and (ii) the histone acetyl transferase in the TGF beta signaling pathway coding *NCOA3* genes, the expressions of both were downregulated in T2D donor islets, and correlated positively with GCG whereas negatively with *INS* (iii), *HMGA1* (variants associated with T2D), and (iv) *BCL2L1* which promotes survival of differentiating pancreatic cells (13), both of which showed higher expression in T2D donor islets and correlated negatively with GCG whereas positively with *INS* (Tables S1, S6, S7).

Several DEGs had previously been shown to be enriched in exocrine cells; of them, seven genes including *MYC, FST* and *SLC38A5* in acinar cells and 17 genes including *KCNJ16* and *CHST15* in ductal cells. The expression of these genes was also negatively correlated with purity, supporting the view of exocrine origin (Supplementary Table 12). Among these DEGs, 29 genes showed higher expression in pancreatic stellate cells compared to endothelial cells (Supplementary Table 13) including *SERPINE2, PTGDS* and *PIEZO2*.

A pathway analysis of the DEGs showed an array of interactions between the various islet cell types involved in T2D pathogenesis (Figure 2iv) including genes enriched in endocrine and exocrine cells (Figure 2iv).

### Functional assessment of two DEGs (*UNC5D and SERPINE2*) by knockdown in human EndoC-βH1 cells

Based on findings in IGW, we set out to functionally validate selected findings.

Expression of ***UNC5D*** was significantly downregulated in islets from T2D donors and negatively correlated with HbA1c levels (Figure 4i, ii). Expression of *UNC5D* correlated with that of *IAPP* (rho = 0.57, p_emp = 0.047) and with *GCG* (rho = 0.36, p = 2.66 × 10^−07^) expression (Supplementary figure 5). The downregulation of *UNC5D* expression in T2D donor islets was also reported in a previous study (7). Immunohistochemical analysis confirmed that UNC5D was expressed in islet cells, and its expression was reduced in beta-cells from T2D donors (Figure 4iii). Moreover, scRNAseq from human pancreatic islets from our extended dataset showed that *UNC5D* is predominantly expressed in beta and deltacells (Figure 4D).

**Figure 4.**
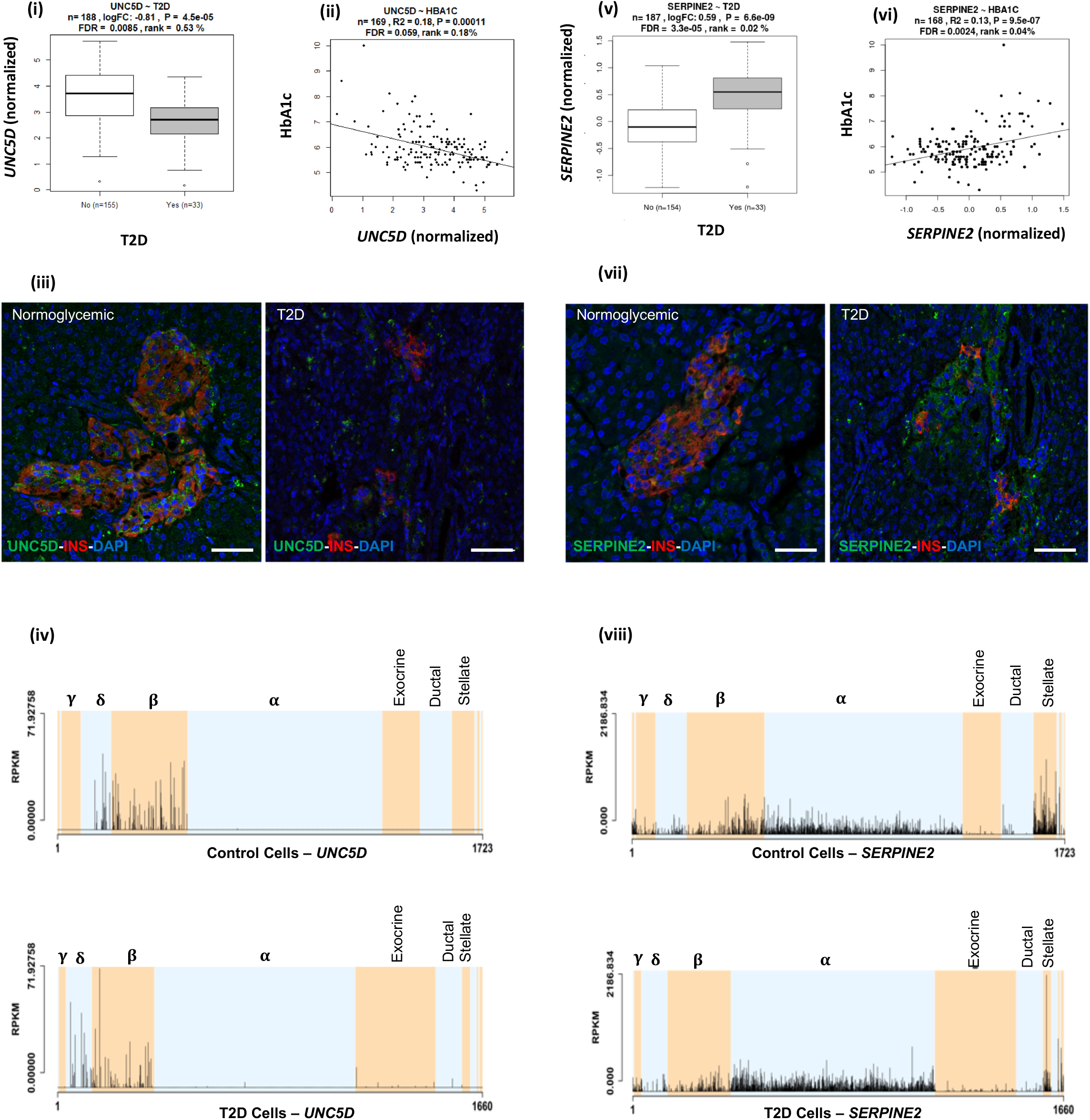
*SERPINE2* and *UNC5D. UNC5D* expression (i) correlated negatively with HbA1c levels and (ii) was downregulated in T2D donor islets. (iii) Immunohistochemical stainings of adult pancreas sections from normoglycemic and type 2 diabetic donors showed *UNC5D* (green) and insulin (red). Scale bar indicate 50 μm, pictures were taken with a 20x objective. Nuclei are shown in blue (DAPI). (iv) in scRNA data from the islets, UNC5D showed expression in delta and beta cells only. (v) *SERPINE2* expression was positively correlated with that of HbA1c levels and (F) upregulated in T2D donor islets. (C) Immunohistochemical stainings of adult pancreas sections from normoglycemic and type 2 diabetic donors showed SERPINE2 (green) and insulin (red). SERPINE2 expression was much higher in immunohistochemistry of the sections from T2D donors Scale bar indicate 50 μm, pictures were taken with a 20x objective. Nuclei are shown in blue (DAPI). (D) In ScRNA data, *SERPINE2* showed ubiquitous expression, with enrichment in pancreatic stellate cells. Expression in alpha cells from T2D donors was significantly higher whereas that in beta cells was lower (although not statistically significant).

Expression of ***SERPINE2*** was strongly upregulated in T2D donor islets, and its expression was strongly positively correlated with HbA1c level (Figure 4v, vi). *SERPINE2* expression was nominally correlated with that of *INS* (rho = 0.17, p = 0.02) and *IAPP* (rho = −0.17, p = 0.019) (Supplementary figure 6). Immunostaining of SERPINE2 in adult human pancreatic sections of normoglycemic donors showed very faint whereas strong islet-associated SERPINE2 immunoreactivity was observed in pancreatic sections of T2D donors (Figure 4vii). Furthermore, scRNAseq data (Martinez-Lopez et al, unpublished) revealed that *SERPINE2* was highly expressed in stellate cells and showed altered expression mainly in alpha cells from T2D donors compared to controls (cZ = 3.85) (Figure 4viii).

We next assessed the functional role of *UNC5D* and *SERPINE2* using small interfering RNA (siRNA) in the well-characterized human β-cell line model EndoC-βH1 human β-cell line (14). By using our previously reported reverse transfection protocol which gives >90% transfection efficiency in this cell line, we achieved a 93±6% (SERPINE2) and 83±4% (UNC5D) mRNA knock-down (KD) upon siRNA transfection (Figure 5A-B) (15). KD of either *SERPINE2* or *UNC5D* had no significant effect on total insulin content (Figure 5C). However, KD of *SERPINE2* (p = 0.001) and *UNC5D* (p = 0.03) significantly reduced GSIS (Figure 5D) but lacked an effect on basal insulin secretion. Moreover, the stimulatory index to glucose (20G/1G) was also significantly reduced in both, by 50±14% (p = 0.001) in *SERPINE2* KD cells and by 64±014% (p = 0.0003) in *UNC5D* KD human beta-cells (Figure 5E). Furthermore, we also determined insulin secretion evoked by 0.5 mM 3-isobutyl-1-methylxanthine (IBMX), a phosphodiesterase inhibitor (16). Interestingly, IBMX-induced insulin secretion was significantly decreased by 35±6.7% (p = 0.0001) in *SERPINE2* KD beta-cells whereas we did not observe changes in *UNC5D* KD beta-cells (Figure 5F). Finally, we studied the impact of KD of *SERPINE2* or *UNC5D* on beta-cell survival when exposed to cytokines (IL-1β, TNF-a and IFN-γ) which are implicated in inflammation in T2D (17). We observed a significantly higher rate of cell death in both SERPINE2 (p = 0.02) and in UNC5D (p = 0.02) KD beta-cells assessed by activated caspase-3/7 levels after exposure to the cytokines cocktail (Figure 5G).

**Figure 5.**
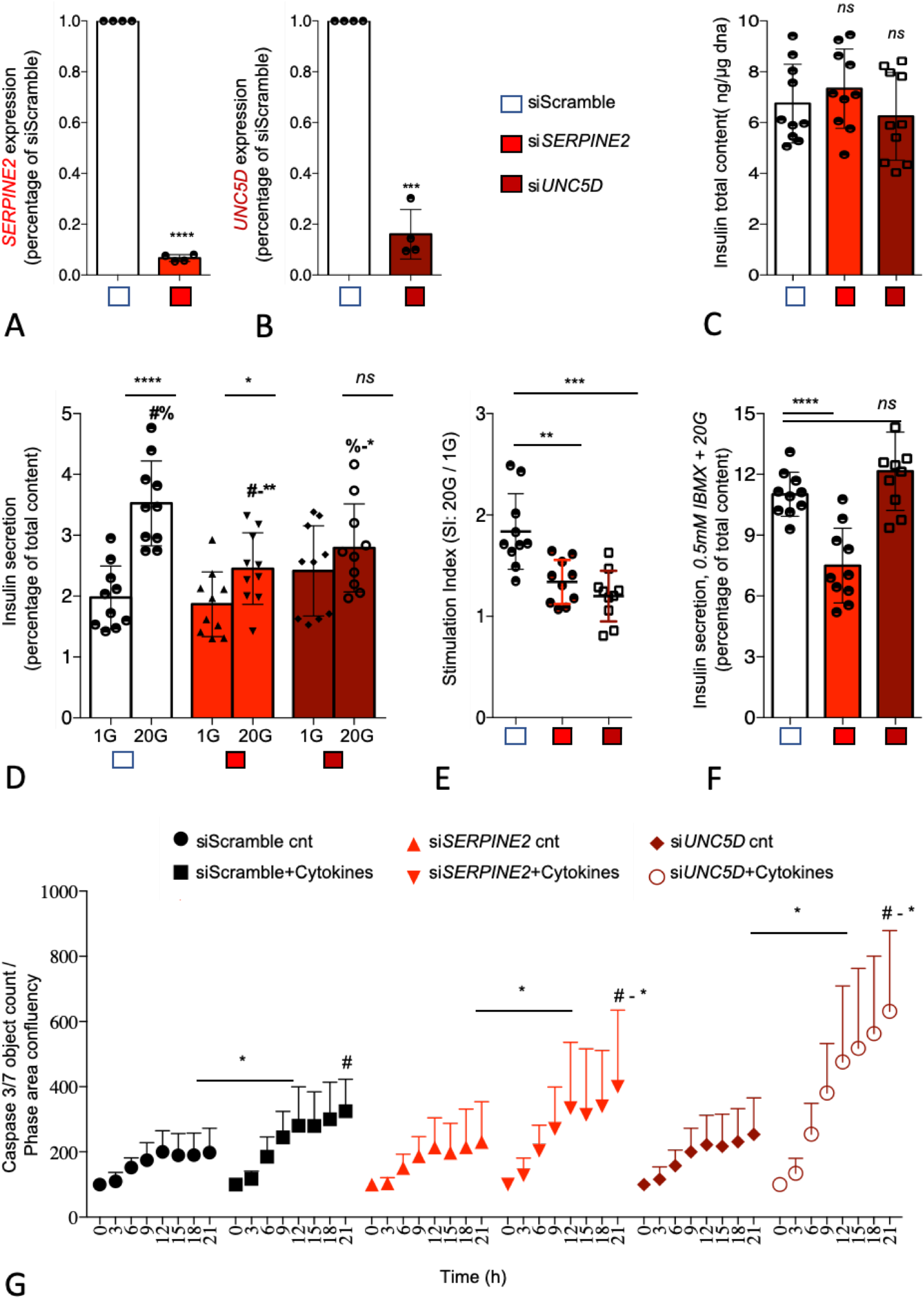
*SERPINE2* and *UNC5D* knock down leads to impaired insulin secretion and induced apoptosis in human pancreatic EndoC-bH1 cells. (**a-b**) Effect of siRNA mediated knock down (KD) on *SERPINE2* and *UNC5D* mRNA. (**c**) intracellular insulin content. (**d**) glucose stimulated insulin secretion. (**e**) stimulation index (ration of high glucose 20G to low glucose 1G). (**f**) insulin secretion stimulated by IBMX. (**g**) Effect of KD of *SERPINE2* and *UNC5D* on cytokines induced apoptosis measured with IncuCyte Caspase 3/7 green reagent. Data are shown as Mean with SD (n = 3 - 5). *p < 0.05, **p < 0.01, ***p < 0.001 (one-way ANOVA, followed by Tukey’s test; #a, b, Two-tailed unpaired t-test, -).

#### Expression quantitative traits (eQTLs)

IGW also provides information on the effect of genetic variation on gene expression in human pancreatic islets. We performed eQTL analysis using GWAS data imputed with the 1000 Genomes Phase I reference panel and RNA data. The effect of SNPs proximal to a given gene on its expression was computed by cis-eQTL analysis using Matrix eQTL. For this analysis, only cis-eQTLs located within 2 Mb of the target genes were considered. Of all 13169 expressed genes with eQTL data, 3243 (24.6%) possessed at least one genomewide significant eQTL (FDR <= 0.05 after adjustment for all tested variants) (Supplementary figure 7A, B). In total, there were 335 343 significant eQTLs.

eQTLs analysis of the DEGs harboured 1948 SNPs associated with expression of 66 genes (Supplementary Table 14). Of particular interest was the Prostaglandin-H2 D-isomerase coding *PTGDS* gene the expression of which was upregulated in islets from T2D donors; it significantly correlated with *INS*, GCG and *IAPP* expression and associated with HbA1c levels (Figure 2, Supplementary figure 7iii). *PTGDS* showed a genome-wide significant eQTL signal from two linked loci which have not previously associated with T2D risk (rs28592848 and rs28375538) (Table S12, Supplementary figure 7iii). Both rs28592848 and rs28375538 showed suggestive signals for association with indices of insulin secretion meaures during IVGTT (acute insulin response, p = 8.65 × 10^−4^ and peak insulin response, p = 7.40 × 10^−4^) (3).

eQTLs at the *IL22RA1, PTGES, CST2, LINC01099, IL1R2, CKAP2, ROR2* and *DHFR* loci exhibited suggestive signals of association with indices of insulin secretion in the MAGIC data analysis (Supplementary Table 15) (18). We recently defined 5 subtypes of diabetes based on commonly measured clinical variables (19). One hundred-twelve eQTLs at 9 gene loci gave rise to suggestive signals of association with the severe insulin deficient diabetes subtype (SIDD), but not with the other subtypes (severe autoimmune – SAID, severe insulin resistant – SIRD, moderately obese – MOD and moderate age related – MARD)(Supplementary Table 16). Of these, the expression of *COTL1* and *ZNRF1* correlated positively with that of *INS* but negatively with that of *IAPP. ZNRF1* expression also correlated negatively with that of *GCG* whereas positively with *SST*. Expression of *CTSF1* also correlated positively with *SST* expression. All three genes were shown to be expressed in beta-cells and were enriched in alpha-cells whereas *PTGER4* was enriched in epsilon-cells (11) (Figure 2D).

### Association betweenT2D genetic risk loci and gene expression in human pancreatic islets

Previous GWAS studies have reported 404 unique variants associated with T2D risk(2). These 404 T2D risk variants nominally influenced the expression of 249 target genes (eGenes). Of the 249 eGenes, 33% were associated with INS expression (empirical P<=0.05), 23% were associated with IAPP expression, 10% were associated with GCG expression and 7% were associated with SST expression. In total, 43% of the eGenes were associated with the expression of one or more genes encoding the islet hormones. Of them, 45 (of the 249 genes) (18%) were found to be associated with T2D (FDR<=0.05) (Supplementary Table 17).

## Discussion

We have harnessed the potential of transcriptome sequencing and genome-wide genotyping in human pancreatic islets to create IGW, a unique research tool that allows the effective integration of genetic information and islet function. To this end, we used islets from 188 donors and combined *in vivo* and *in vitro* functional studies and obtained novel insights into molecular mechanisms underlying dysregulation of glucose metabolism and impaired islet function in T2D. We explored how gene expression was altered in islets from T2D donors and characterized the differentially expressed genes (DEGs), e.g., by studying their expression pattern in other tissues (fat, liver and muscle) from the same donors, and their co-expression with islet hormones. We also examined genetic regulation of gene expression and their link to islet function. *SERPINE2* (upregulated in T2D) and *UNC5D* (downregulated in T2D) genes were selected for functional analysis to explore their potential role in T2D pathogenesis. Finally, we present Islet GeneView, a web resource to visualize information on genes of interest in this comprehensive catalogue of gene expression in islets.

IGW is complementary to other related databases such as the Islet eQTL explorer (20), and TIGER (http://tiger.bsc.es/), which connect genetic variation to expression and chromatin state, and GTEx (5) which comprehensively characterizes the human transcriptome across many tissues, including whole pancreas. GTEx, however, does not provide data on the islet transcriptome, which clearly is very different from that of the total pancreas which includes the exocrine part of the pancreas. Several studies have focused on human pancreatic islets to obtain insights into general molecular mechanisms of diabetes (8, 20, 21),(22). Our study provides another perspective by (a) exploring and characterizing DEGs and (b) by providing a web resource for visualization of gene expression which can be linked to glycemic status, expression of islet hormones and other relevant phenotypes.

IGW identified 285 genes whose expression was altered in islets from T2D donors. Of them, changes in expression of 39 genes were also observed in at least one other dataset (7) using microarray data from organ donors and partially pancreatectomized individuals. Nine genes overlapped between all three datasets with directional consistency. This variability in the transcriptomes across the datasets could partially be explained by the differences in protocols for obtaining and processing the islets, as well as by the method of transcriptome profiling. However, it could also be attributed to other potential sources of variation, not least, differences in proportion of endocrine component (“purity”) as well as study power and population differences. In addition, the now well recognized variability in T2D phenotypes (19) may also contribute sample variability in the different studies.

Among the replicated genes were *CHL1, SLC2A2* and *HHATL*, which were also reported in our previous studies based on a subset of the current islet dataset (8). *CHL1* has also previously been shown to be associated with T2D (23). Our data suggest that high expression of neural adhesion molecule L1 encoding *CHL1* in islets could have beneficial effects on beta-cell function. In support of this, *CHL1* expression was downregulated in islets from T2D donors and correlated with insulin content (24). In addition, an eQTL for *CHL1* was associated with fasting insulin concentrations (9, 10). KD of *CHL1* in a clonal rat insulinoma cell line (INS1) resulted in impaired GSIS (24). *SLC2A2* encoding the low affinity/high capacity GLUT2 glucose transporter, has previously been shown to play a key role in islet function (25) and common variants in the *SLC2A2* gene are associated with T2D and insulin secretion. Variants in this gene have recently been shown to alter glycemic response to metformin therapy in recently diagnosed T2D patients (26). Indeed, it has been shown that low birth weight in individuals with *SLC2A2* deficiency have lower insulin secretion, This reduction in insulin may play a key role in fetal development, particularly in the third trimester, given the growth-promoting effects of insulin (27). The expression of all these three genes correlated with insulin secretion in islets as well as with islet hormone expression. Of relevance to insulin secretion was also the gene for glycine receptor subunit alpha 1 (*GLRA1*), the expression of which was down-regulated in T2D islets, and significantly correlated with stimulatory index. *GLRA1* has been suggested to facilitate insulin secretion through a glycine-insulin autocrine feedback loop (28). Of note, GLRA1 undergoes DNA methylation after exposure of human islets to high glucose concentration and its KD in INS-1 cells blunts GSIS (29). Moreover, *GLRA1* and *HHATL* showed higher expression in beta-cells. Taken together, the data imply that these genes could serve as potential therapeutic targets in T2D.

The DEGs in IGW showed heterogeneity in expression between different islet cell populations, both endocrine and exocrine. These included genes enriched in (i) beta-cells (and associated with insulin secretion e.g. *GLRA1* and *HHATL*), (ii) other endocrine cell types (alpha-cells-e.g. *PPP1R1A*, gamma-cells – e. g. *INPP5F*, delta-cells – e.g. *OPRD1* and e.g. *FFAR4*, epsilon-cells – e.g. *PTGER4*) (iii) exocrine cells (acinar cells – e.g. *MYC*, ductal cells – *CDKN1A*) (iv) pancreatic stellate cells – *SERPINE2*. Further, co-expression studies might help to identify clusters of genes sharing common mechanisms for regulation. For instance beta-cell enriched genes co-expressed with *INS* or *IAPP* as well as alpha-cell enriched genes co-expressed with *GCG* could potentially indicate a key role in hormone secretion and thereby represent downsteam targets. Through pathway analysis using a protein-protein interaction database as reference, we could show interaction between genes enriched both endocrine and exocrine cells. These phenomena could be interpreted as interactions of endocrine-exocrine components regulating islet function.

Genes identified by IGW showing tissue-specific expression could play a critical role in function of the specific tissues. Identification of such genes has provided molecular insights into tissue function and disease pathogenesis. To this end we discovered DEGs that showed islet-specificity and islet-enrichment in expression compared to fat, liver and muscle from the same individual; some of these were beta-cell enriched (*GLRA1, SSTR5-AS1, IAPP, CAPN13, GLP1R, DACH2* and *LINC01099*), while others were alpha-cell enriched (*CRH, SYNDIG1, CALY, KCNC1*, and others). A vast majority of the DEGs showed expression in all the tissues tested implying their pleiotropic nature (associated with multiple phenotypes associated) (22, 30). Tissue specificity and enrichment information are vital when evaluating novel therapeutic targets.

We focused more in depth on two interesting differentially expressed candidate genes, *UNC5D* and *SERPINE2* by exploring them further in the human beta-cell line EndoC-βH1. UNC5D is the newest member of Uncoordinated-5 (*UNC-5*) receptor family which act as receptor for axon guidance factors, Netrins. Netrin factors are well described to be important for beta-cell development and survival (31, 32). However, the functional role of *UNC5D* in adult human beta-cells remains largely unknown. *UNC5D* is predominantly expressed in beta and delta-cells of human islets and importantly, is one of the top downregulated genes in T2D donor islets (present dataset and scRNAseq data). Notably, none of the other genes of the UNC5 family receptors showed any differential expression pattern while Netrin (*NTN1*) showed only a marginally reduced expression in the T2D donor islets (p = 0.01). Further, we observed that *UNC5D* gene expression also showed strong positive correlation with two known T2D genes *ROBO2* and *SLC30A8* (Supplementary figure 8). It was interesting to see the strong association of *UNC5D* with the T2D target gene, *ROBO2*, which is known to play a role in beta-cell survival under different stress conditions through Slit-Robo signaling (33). *UNC5D* expression is unique to human islets with no expression in the mouse islets (34). Likewise, mouse islets show a unique expression of *Unc5a*, unlike in human islets, suggestive of species-specific activity of these receptors (Supplementary figure 9). Our observations warrant an in-depth study to gain more insight into the effect of *UNC5D* on insulin secretion and beta-cell survival, possibly through a Netrin-UNC5; Slit-Robo signalling loop.

*SERPINE2* (Serpin Peptidase Inhibitor, Clade E, Member 2), also known as Protease Nexin-1 (PN-1) is another interesting DEG the expression of which was strongly upregulated in the T2D donor islets. Notably scRNAseq showed that *SERPINE2* is predominantly expressed in alpha and stellate cells, and increased in T2D islets. However, its expression in beta-cells was low and marginally changed in T2D islets (Figure 4H). Moreover, *SERPINE2* expression has previously been reported to be downregulated in the microvascular endothelium of patients with diabetic neuropathy through the upregulation of specific microRNAs (35). We employed RNA interference to mediate loss of function to understand the role of *SERPINE2* in human beta-cells. Depletion of *SERPINE2* mRNA resulted in a markedly decreased insulin secretion in response to glucose and IBMX. *SERPINE2* is an extracellular matrix (ECM) remodeler glycoprotein that acts as an inhibitor for trypsin, thrombin, and urokinase plasminogen activator like serine proteases (36). *SERPINE2* may potentially affect beta-cell function and survival through its actions on the extracellular milieu but more detailed studies are needed to verify this.

GWAS have reported 403 genetic loci associated with T2D risk although the mechanisms remain largely unknown. To gain insight into mechanistic effects of SNPs, we explored whether they influence expression of unique genes (eQTLs). To this end, we identified multiple gene loci influencing gene expression in islets; the expression of these genes was altered in islets from T2D donors and correlated with insulin secretion. *PTGDS* was one such gene; it encodes an enzyme converting prostaglandin H2 to prostaglandin D2 (PGD2). High PGD2 levels have been shown to enhance insulin sensitivity (37), and KD of *PTGDS* results in an opposite effect resulting in insulin resistance, but also nephropathy and atherosclerosis. While adipose tissue could play a role here, insulin resistance has in many studies been shown to play a key role in pathogenesis of nephropathy. Interestingly, nephropathy was primarily seen in the insulin-resistant subgroup of SIRD in the new classification of T2D (19). While we found little support for the correlation between diabetes and *PTGDS* in islet cells in previous studies, the current results suggest that expression of the gene in islets is correlated both with diabetes status, BMI and *INS* expression, indicating that *PTGDS* function and prostaglandin levels may be connected to insulin secretion as well as peripheral insulin resistance.

Most of the results reported here can be found in the Islet Gene View. This is a web application that we have developed to visualize gene expression in simple histograms. In addition, information on purity allows for a partial separation of expression patterns in endocrine and exocrine tissue, as strong positive correlation of expression of a gene on purity is indicative of a high proportion of exocrine tissue. Relationship to glycemic status, BMI and related phenotypes is provided as simple-to-read graphs for a specific gene.

A strength of IGW is the large sample size of islets from organ donors with maintained blood circulation, which is larger (+99 donors) than previous publications from our centre (8, 24). Following a previous report on a smaller subset (8), we have made several refinements to the analytical pipeline including batch correction using ComBat. We have applied refined methodology for calculating p-values for correlations, which are now independent of batch effects. This reduces the influence of nonspecific inter-gene correlations resulting from the normalization procedure for gene expression. All gene-gene correlations have been precalculated in order to estimate the null distribution of the correlation values. This is computationally intensive, but only has to be done once. One limitation is that the graphs and data are descriptive and cannot distinguish correlation from causation. However, we selected two genes for functional validation, demonstrating that the data can be used to further explore functionally relevant genes expressed in islets.

Taken together Islet Gene View is a tool to facilitate research on human pancreatic islets and will be made accessible to the entire scientific community. The exploratory use of IGW could help designing more comprehensive functional follow-up studies and serve to identify therapeutic targets in T2D.

## Materials and methods

### Sample acquisition

Human pancreatic islets (n = 188), fat (n = 12), liver (n = 12) and muscle (n = 12) were obtained through the EXODIAB network from the Nordic Network for Clinical Islet Transplantation (http://www.nordicislets.org). All procedures were approved by the ethics committee at Lund University. The isolation of total RNA including miRNA was carried out using the miRNeasy (Qiagen) or the AllPrep DNA/RNA (Qiagen) mini kits as described previously (8). The quality of isolated RNA was controlled using a 2100 Bioanalyzer (Agilent Technologies) or a 2200 Tapestation (Agilent Technologies) and quantity was measured using NanoDrop 1000 (NanoDrop Technologies) or a Qubit 2.0 Fluorometer (Life Technologies). Clinical characteristics of the donors are shown in Supplementary Table 1.

### Islet Phenotypes

Purity of islets was assessed by dithizione staining, and estimates of the contribution of exocrine and endocrine tissue was assessed as previously described (38).

#### Phenotypic information

Diagnosis of Type 2 diabetes (T2D) was either based on a clinical diagnosis of T2D (N = 33) or on an HbA1c above 6.5% (NGSP units; equal to 48 mmol/mol in IFCC) (N = 25). IGT was defined as HbA1c between 6 and 6.5% (N = 30).

Information on gender and BMI was obtained from donor records. Stimulatory index (SI) was used as a measure of GSIS. For this purpose, islets were subjected to dynamic perfusion of glucose, which was raised from 1.67 to 16.7 mmol/l for one hour; insulin was measured at both high and low glucose. The fold change in insulin levels between the two conditions was used as a measurement of glucose-stimulated insulin secretion.

### Sample preparation for RNA sequencing

One μg of total RNA of high quality (RIN>8) was used for sequencing with a TruSeq RNA sample preparation kit (Illumina). We here included 99 islet samples in addition to the 89 islet samples and processed them uniformly following the same protocol as described previously (8). Briefly, the size selection was made by Agencourt AMPure XP beads (Beckman Coulter) aiming at a fragment size over 300 bp. The resulting libraries were quality controlled on a 2100 Bioanalyzer and a 2200 Tapestation (Agilent Technologies) before combining 6 samples into one flow cell for sequencing on a HiSeq 2000 sequencer (Illumina).

### Islet Gene View

#### RNAseq Data analysis

The raw data were base-called and de-multiplexed using CASAVA 1.8.2 (Illumina) before alignment to hg38 with STAR version 2.4.1. To count the number of reads aligned to specific transcripts, featureCounts (v 1.4.4) (39) was used, with GENCODE version 22 as gene, transcript and exon models. The average number of counts mapped to genes was 26.1 million reads (± 13.1 million) ranging from 10.0 million up to 79.4 million.

Raw data were normalized using trimmed mean of M-values (TMM) implemented in edgeR and transformed into log2 counts per million (log2 CPM), using voom (40). Samples with less than 10 million reads in total were excluded from further analysis. In addition, only genes with more than 0 FPKM in 95% of samples and an average expression of more than 1 FPKM were retained, leaving 14,108 genes for analysis in the 188 samples.

A potential association between gene expression and phenotypes was analyzed by linear modeling. Voom was used to calculate variance weights, linear modeling was performed with lmFit, and P-values were calculated using the eBayes function in limma (41). P-values adjusted for multiple testing were calculated across all genes using Benjamini-Hochberg correction (42).

As expression of 50% of the genes in the dataset was correlated with purity (mostly due to admixture of exocrine tissue), we included purity as a covariate in the linear models for all association analyses, together with sex and age. Six individuals did not have T2D based upon a clinical diagnosis but based upon HbA1c above 6.5% and were excluded from the differential expression analysis of T2D versus controls.

An empirical and conservative approach was used to calculate P-values for gene-gene correlations. 1 million gene/gene pairs were randomly selected from all genes after filtering for expression and the Spearman correlations for the pairs were calculated. This provided a background distribution of gene expression correlation. To calculate P-values from this background distribution for a given correlation, the proportion of background gene pairs with the same or more extreme correlation values was used. This provides a more robust method for detecting gene-gene pairs with high co-expression.

To investigate the biological relevance of the RNA-Seq and eQTL data, we investigated the relationship between previously known risk variants and gene expression in Islet Gene View (IGW). For a list of 404 variants previously associated with either T2D risk (2) or with β-cell function, we identified all variants nominally significant as eQTLs in the islets. The list of genes affected by eQTLs was tested for enrichment of genes whose expression was correlated with secretory genes using bootstrapping. This was done by repeatedly selecting random sets of genes of the same size (10 000 iterations). The mean empirical P-value for each run was then calculated and was used as a null distribution to calculate the probability of the observed mean P-value in the observed set of genes.

Pathway analysis: For the 285 genes which were differentially expressed between islets from T2D compared to normoglycemic donors, pathway analysis was performed using the InWeb(43) protein-protein interaction network implemented in GeNets (44). GeNets visualization of network data was accessed at http://apps.broadinstitute.org/genets and Intomics (https://inbio-discover.intomics.com/map.html#search).

##### Genome-wide genotying and imputation

Genotyping was performed using an Illumina OmniExpress microarray and QC was performed as previously described (8, 22). The resulting genotype data was phased using the SHAPEIT version 2 software and imputed with IMPUTE2, using the 1000 genomes phase 1 integrated variant set as described previously.

##### eQTL analysis

eQTL analysis was performed using gene expression data normalized for batch effects with ComBat, and the directly genotyped, as well as the imputed, genotype data. Only cis-eQTLs within 2 MB of the target gene were considered. The R package MatrixeQTL was used to analyze the association between gene expression and genetic variants.

##### Islet geneview web application

A website to access the resulting plots was developed using the Shiny web framework. A table of available genes can be searched and used to select a gene to investigate, which shows the corresponding set of plots for that particular gene. These plots were generated using an R script.

#### Immunohistochemistry

Human pancreatic islets were processed for paraffin (6 μm sections) and cryo-embedding (10 μm sections) respectively. Immunostaining was performed as described previously(45) with the following antibodies: guinea pig α-insulin (1:2000, Millipore/1:800, DAKO), mouse α-glucagon (1:2000, SIGMA), mouse SERPINE2 (1:2000, (LSBio Catalog # LS-C173926), and goat UNC5D (1:500, (Novus Bio, Catalog# AF1429). Nuclear counterstaining was performed using 4,6-diamidino-2-phenylindole (DAPI,1:6000, Invitrogen).

### EndoC-βH1 cells

Human β-cell line EndoC-βH1 was obtained from Univercell Biosolution S.A.S., France (1). The cells were cultured on Matrigel (1%) and fibronectin (2 μg/ml) (Sigma-Aldrich) coated plates in low-glucose (1g/L) Dulbecco’s modified Eagle’s medium (DMEM; Invitrogen) at 37°C and 5% CO2 as previously described (14).

### siRNA Transfection

EndoC-βH1 cells were transfected using Lipofectamin RNAiMAX (life technologies) and 30nM ON-TARGET *plus* siRNA SMARTpool for human *SERPINE2* (Horizon; L-012737-00-0005) or *UNC5D* gene (Horizon; L-015286—00-0005) or 20 nM ON-TARGET *plus* Nontargeting pool (siNT or Scramble) (Horizon; D-001810-10-05) as described (15). Cells were harvested 96 h post-transfection for further studies.

### Quantitative RT-qPCR

Total RNA was extracted from EndoC-βH1cells using Macherey-Nagel RNA isolation kit. cDNA was prepared using the Maxima first stand cDNA synthesis kit as per manufacturers recommendations (Thermo Fisher scientific). Briefly, 5x Hot FIREPol EvaGreen qPCR mix plus for quantitative PCR (Solis Biodyne) was used for the reactions with a Corbett Rotor-Gene 6000 (Qiagen). The reactions were pipetted with a liquid handling system (Corbett CAS-1200, Qiagen). All reactions were performed in duplicates on at least three biological replicates. Cyclophilin-A was used as an endogenous control. Primer sequences are available upon request.

### Insulin secretion and content

EndoC-βH1 cells were transfected with siRNA on matrigel and fibronectin coated 24well plates at 2 × 10^5^ cells per well. Following 96h of siRNA transfection cells were incubated in 1mM glucose containing EndoC-βH1 culture medium for over-night and next 60 mins in βKREBS (Univercell Biosolution S.A.S., France) without glucose. Cells were sequentially stimulated with 500 μl βKREBS containing 1mM glucose, 20 glucose and then finally 20mM glucose + 0.5 mM IBMX (#-Isobutyl-1-methylxanthine; #I5879, Sigma-Aldrich) each for 30 mins at 37°C in a CO_2_ incubator. After every incubation the top 250 μl supernatant was carefully collected; at the end of the experiment cells were washed and lysed with TETG (Tris pH8, Trito X-100, Glycerol, NaCl and EGTA) solution prepared as per Univercell Biosolution EndoC-βH1 manual guide for the measurement of total content. Secreted insulin after each stimulation was added to the final content to determine the % of secretion of the total content. For the measurement of total insulin content 96h post-transfection cells were washed twice with PBS and lysed with TETG and stored in −80°C until insulin ELISA measurement. Simultaneously DNA content was also determined to normalized the total insulin content. Secreted and intracellular insulin were measured using a commercial human insulin ELISA kit from Mercodia (Sweden) as per manufacturer’s instruction.

#### Caspase-3/7 Apoptosis assays

EndoC-βH1 cells, 2 × 10^5^ cells per well of 24-well plate (Costar #3526) were treated with siRNA against *SERPINE2 or UNC5D* or Non-targeted control as described previously in siRNA methods section. After 96 h siRNA treatment, cells were washed twice with PBS and stimulated with cytokine cocktail consisting of IL-1β (5 ng/ml, R&D Systems, Minneapolis, MN, USA), and IFN-γ (50 ng/ml, R&D Systems) and TNF-α (10 ng/ml, R&D Systems) in the presence of IncuCyte Caspase-3/7 green reagent (IncuCyte, Essen Bioscience, #4440, 1:2000) for next 24 h. Every 3h images were taken with an IncuCyte-S3 Live-Cell Imaging system (Essen Bioscience) using 488 nm laser. Images were analyzed with IncuCyte S3 software and apoptosis has been quantified as the ratio of green fluorescent Caspase-3/7 green active object count to phase area confluency.

### Lookups

#### Indices of insulin secretion

For this lookup, we selected SNPs which associated with gene expression (eQTLs) of the 285 DEGs. We performed a lookup of these SNPs in the Meta-Analyses of Glucose and Insulin-related traits Consortium (MAGIC) analysis for association with indices of insulin secretion (corrected insulin response – CIR and disposition index (DI) (18).

#### Association with diabetes subtypes

We recently stratified adult individuals with newly diagnosed diabetes from Skåne, Southern Sweden (all new diabetics in Skåne – ANDIS study) into 5 subtypes based on commonly measured clinical variables. These included: glutamate decarboxylase autoantibodies (GADA), body mass index (BMI), glycosylated hemoglobin (HbA1c), age at diabetes onset, gender, beta-cell function (HOMA2-B) and insulin resistance (HOMA2-IR) estimated from fasting glucose and C-peptide (19). The five clusters included: Severe Auto-Immune Diabetes (SAID), Severe Insulin-Deficient Diabetes (SIDD), Severe Insulin-Resistant Diabetes (SIRD), Mild Obesity-related Diabetes (MOD and Mild Age-Related Diabetes (MARD). These clusters were phenotypically different and manifested different outcomes in terms of risk for complications. GWAS was performed on these individuals in an effort to identify cluster specific risk variants, whereby the association with each subtype of diabetes was assessed against controls from the same population. We here looked up the eQTLs for DEG to assess if they showed cluster specific signals, with specific focus on those associated with the insulin deficient SIDD subtype.

#### Cause of consequence

In our previous study(12), we treated human pancreatic islets from normoglycemic donors and hyperglycemic donors with normal and high glucose for 24 hours and analyzed gene expression changes to distinguish between genes which could be causally involved in pathogenesis from those that were likely a consequence. We here looked up genes whose expression was altered in islets from T2D individuals (differentially expressed genes – DEG) in this dataset to assess if they were also impacted by acute glucose exposure, and if they also associated with indices of insulin secretion.

## Acknowledgements

Human pancreatic islets, muscle, fat and liver samples were provided by The Nordic Network for Clinical Islet Transplantation. The work in this paper has been financially supported by grants from the Swedish Research Council: strategic research environment grant (EXODIAB, 2009-1039) and project grant (2015–2558) to LG, networking grant (2015–06722) to RP; collaborative grant from the Swedish Foundation for Strategic Research to the Lund University Diabetes Centre (IRC 15-0067); JDRF(award 31-2008-416); Diabetes Wellness grant to RP (720-858-16JDWG); collaborative grants with Regeneron and Eli Lilly to LG. We thank Mattias Borell, Maria Sterner, Malin Neptin and Malin Svensson for technical support. We also want to express our deepest gratitude to the deceased organ donors and their relatives.

## Author contributions

RPB, OA, PS and VC analysed the data, RPB, PS, OA and LG designed the study. RBP, OA and VC interpreted the results and wrote the manuscript, with additional input from ER, OH, HM, OK and UK. VC, EOL and UK performed laboratory experiments and measurements. RBP supervised the project. LC contributed the data. All authors provided critical feedback and helped shape the research, analysis and manuscript.

## Data availability

All relevant data supporting the key findings of this study are available within the article and its Supplementary Information files or from the corresponding author upon reasonable request. Genotype, technical and biological covariates, and sequence data have been deposited at the European Genome-phenome Archive (EGA; https://www.ebi.ac.uk/ega/) under the following accession numbers: EGAS00001004042 [https://ega-archive.org/studies/EGAS00001004042]; EGAS00001004044 [https://ega-archive.org/studies/EGAS00001004044], EGAS00001004056 [https://ega-archive.org/datasets/EGAS00001004056].

---

**Supplementary figure 1.**
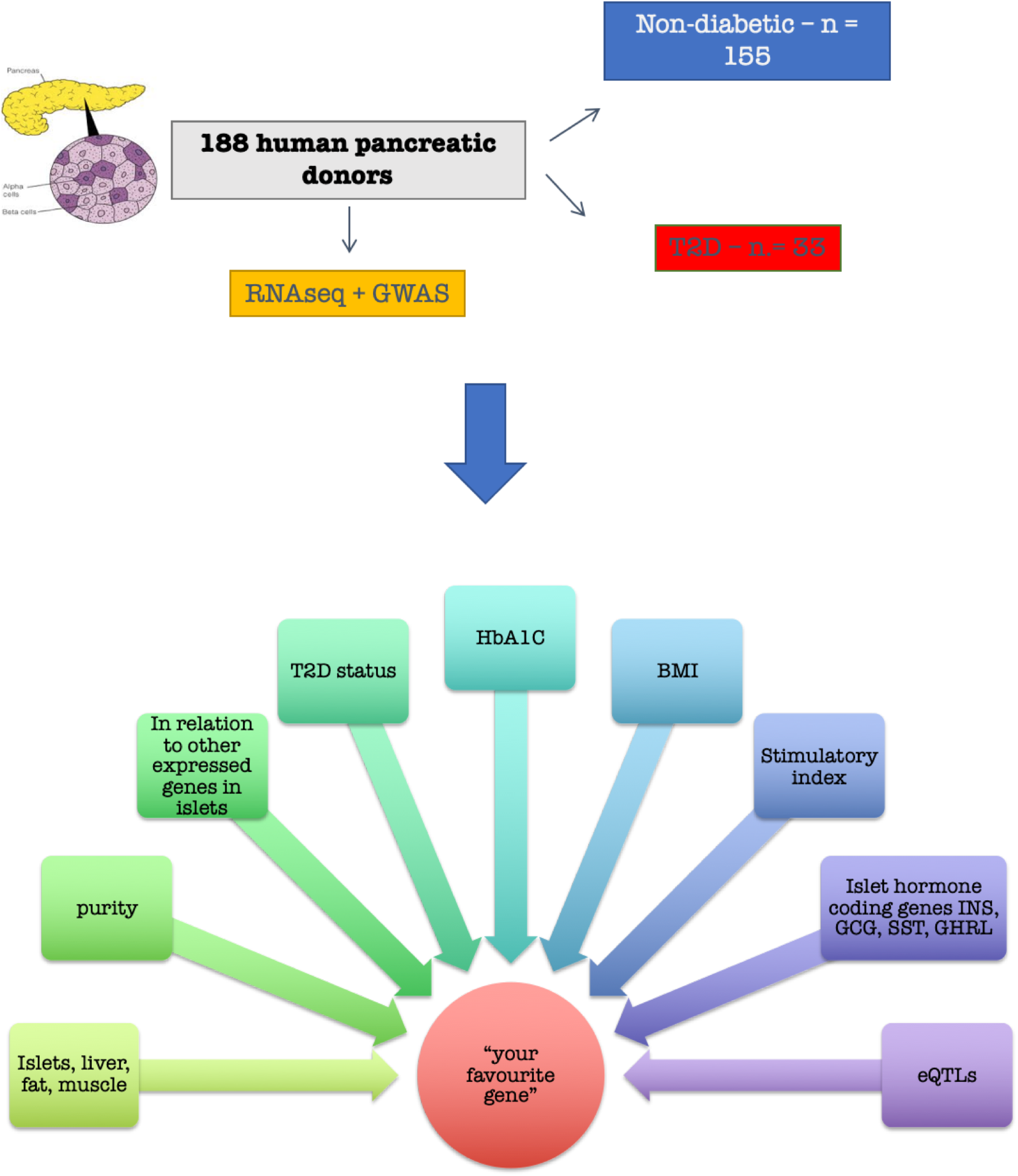
Study design of Islet Gene View (IGW). A: Islets from human organ donors (n = 188, 155 non-diabetic and 33 with type 2 diabetes, were isolated for RNA sequencing. A: expression of the gene in fat, islets, liver, and muscle, in the same pool of 12 individuals. B: Gene expression as a function of purity, defined as the percentage of endocrine tissue. C: Expression of the selected gene in relation to other genes in islets. D-H: Gene expression in relation to several diabetes-related phenotypes, i.e. T2D diagnosis (figure 1D), HbA1c stratum (1E), continuous HbA1c (1F), BMI (1G), and stimulatory index (1H). Test statistics are reported, namely: coefficient of determination (R2), nominal P-value, and percentage rank among all genes as calculated based on sorted P-values. I-M: Gene expression in relation to the islet hormone coding genes; SST, INS, GCG, PPY and GHRL. Spearman’s ρ (r) and the P-value of the gene based on the empirical correlation distribution is reported. INS = Insulin, GCG = glucagon, SST = somatostatin, PPY = pancreatic polypeptide, IAPP = islet amyloid polypeptide.

**Supplementary figure 2.**
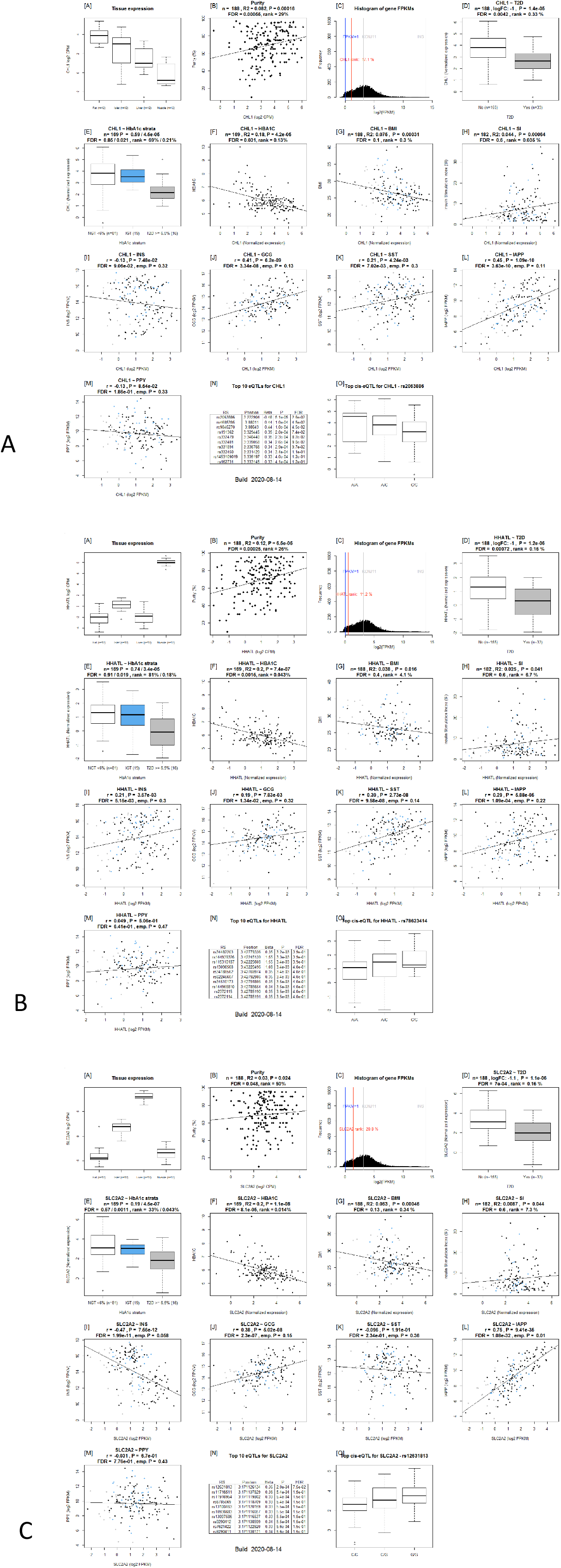
Islet gene view plots for (A) CHL1 (B) HHATL and (3) SLC2A2. The expression of all three genes were downregulated in T2D donor islets, correlated with HbA1c levels, and nominally with BMI, and SI. CHL1 and SLC2A2 correlation negatively correlated with that of INS whereas HHATL was positively correlated. The expression of HHATL correlated with that of INS, SST and IAPP, CHL1 with GCG and SLC2A2 with IAPP

**Supplementary figure 3:**
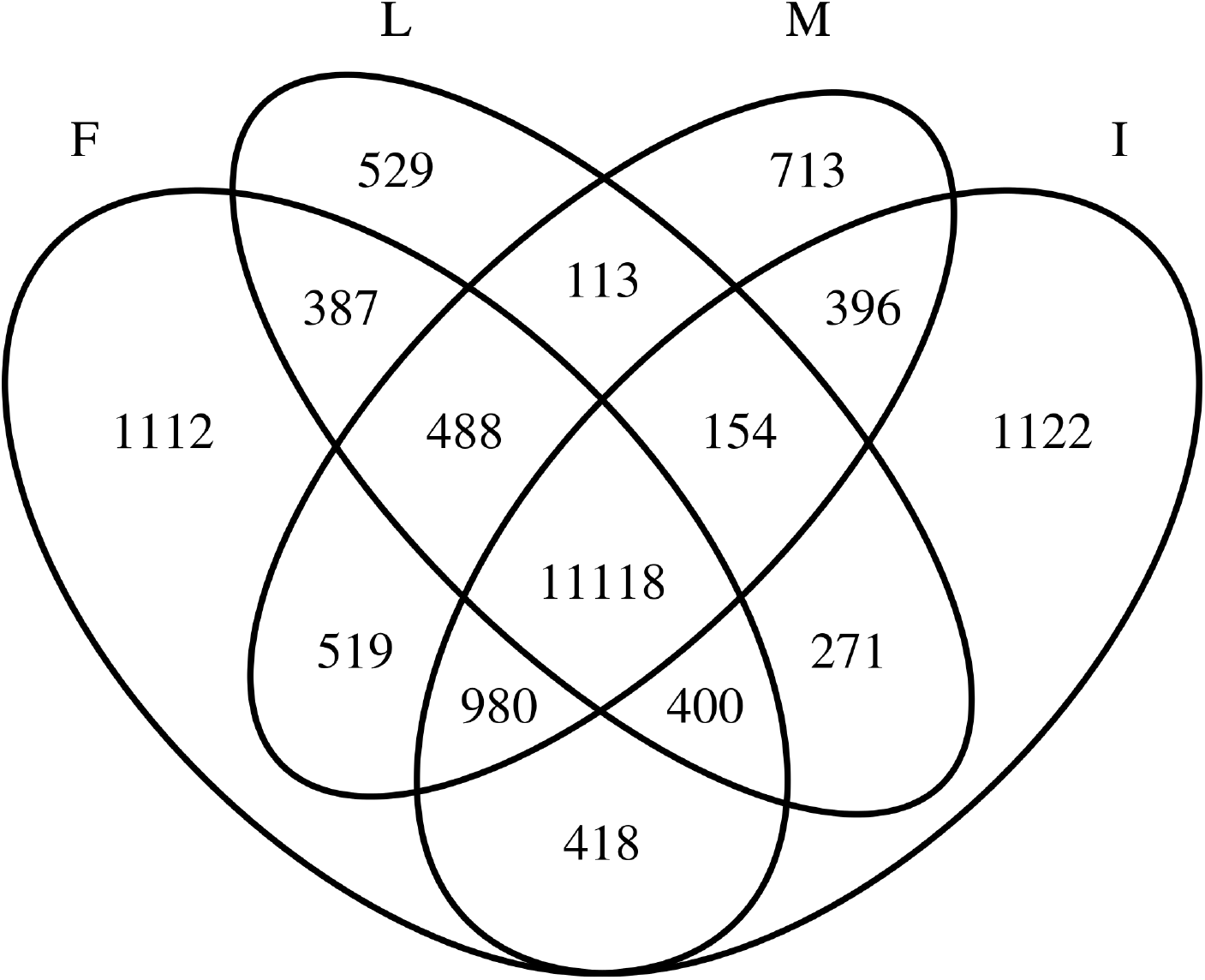
Overlap of genes with expression detected in all 12 samples per tissue in fat(F), liver(L), muscle(M) and islets(I).

**Supplementary figure 4:**
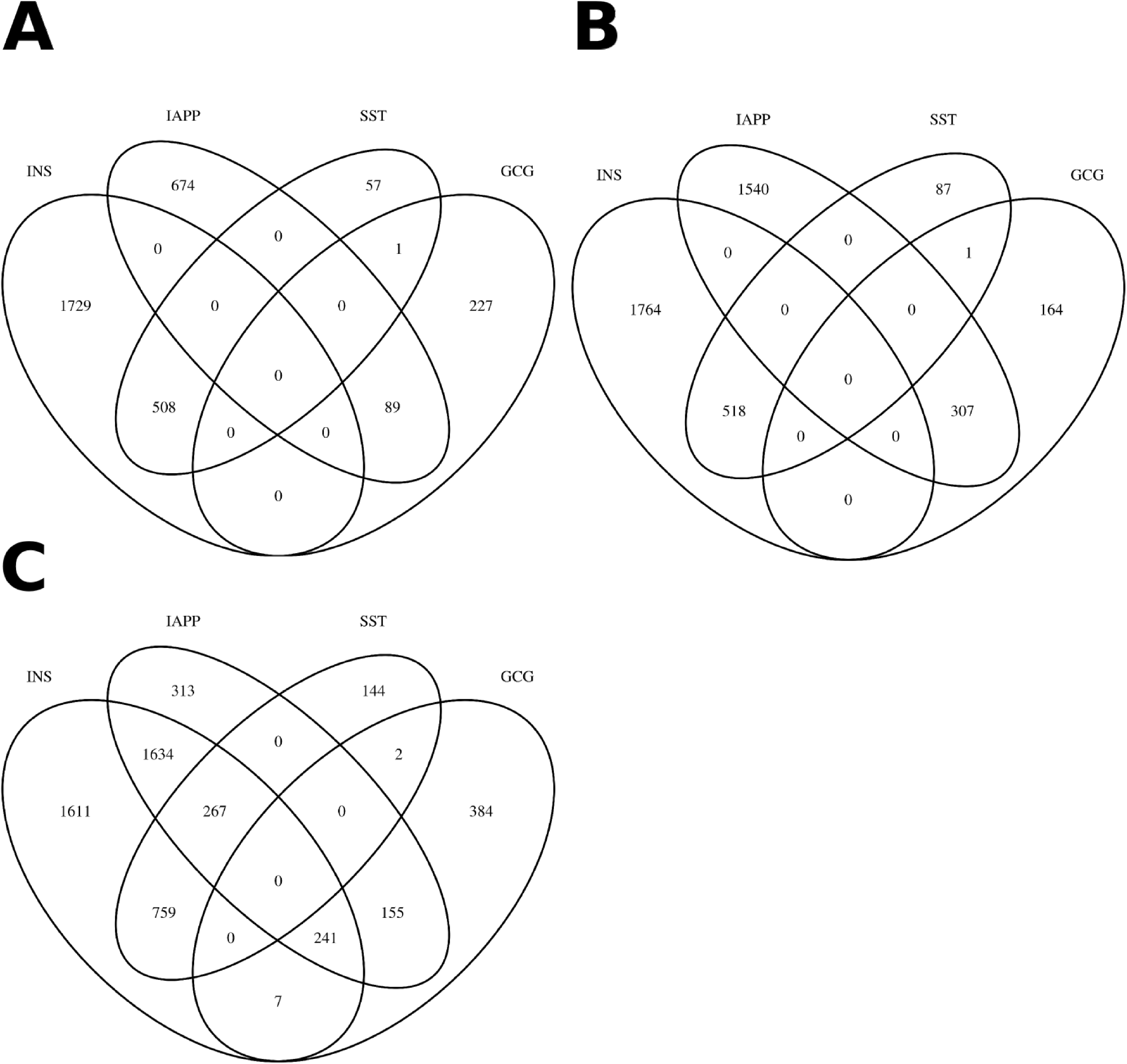
Co-expression with secretory genes Venn diagram of genes co-expressed with secretory genes. A: Overlap of positively co-expressed genes (ρ ≥ 0, P ≤ 0.05) with insulin (INS), glucagon (GCG), somatostatin (SST) and islet amyloid polypeptide (IAPP). B: Overlap of negatively co-expressed genes (ρ ≤ 0, P ≤ 0.05). C: overlap of all co-expressed genes regardless of direction (P ≤ 0.05).

**Supplementary figure 5:**
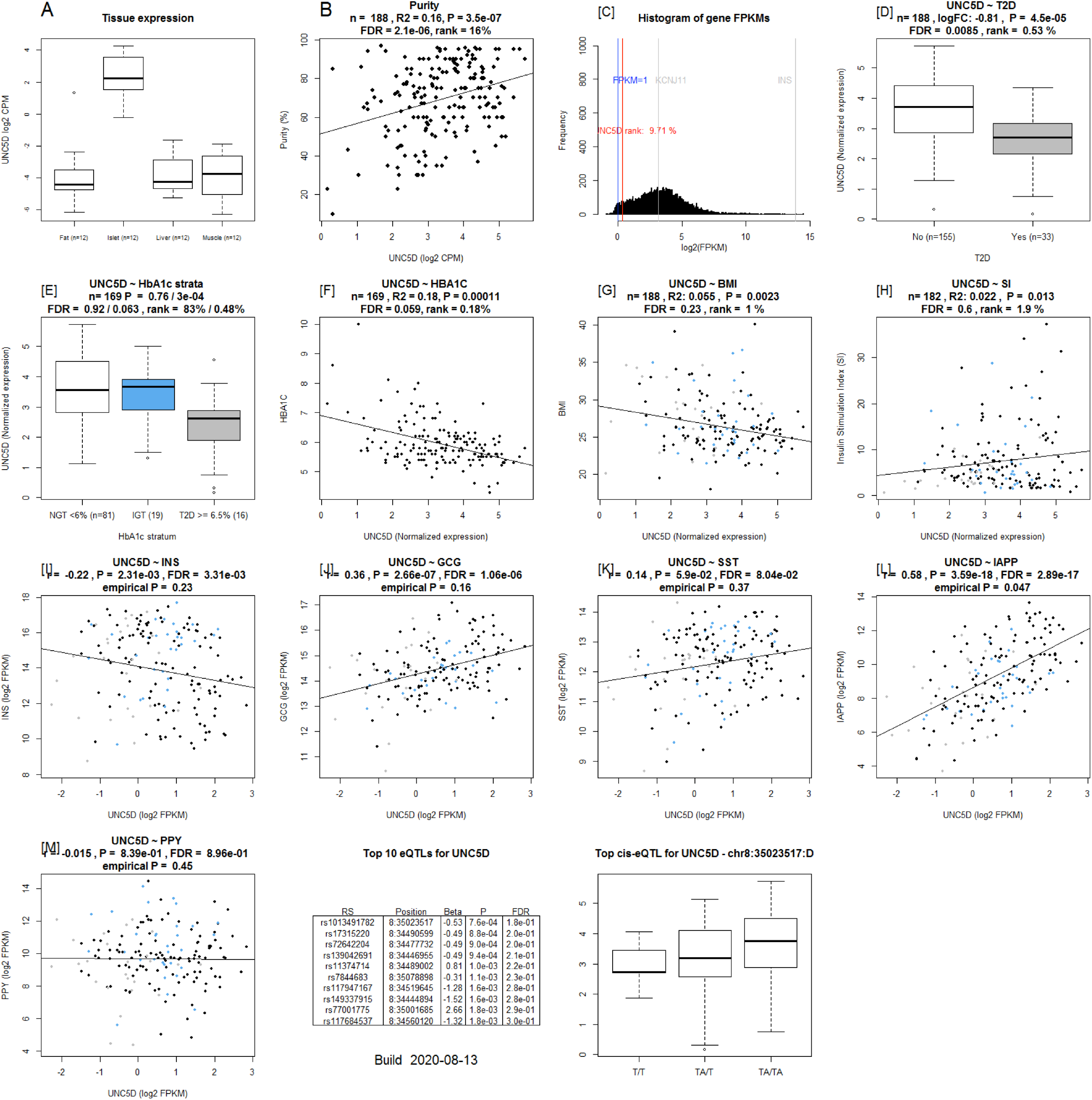
Islet Gene View graph of *UNC5D. UNC5D* shows highest expression in islets compared to in fat, liver, and muscle in the same pool of 12 individuals [A], UNC5D is positively correlated with purity, (endocrine component) [B] and is ranked within the top 9.71% of genes expressed in the islets [C]. UNC5D expression is downregulated in T2D donor islets compared to controls [D], as well as in IGT and T2D donors compared to NGT [E] and is negatively correlated with HbA1c levels [F] UNC5D expression in nominally negatively correlated with BMI [G] and positively with stimulatory index [H]. Test statistics are reported, namely: coefficient of determination (R2), nominal P-value, and percentage rank among all genes as calculated based on sorted P-values. [I-M]: Gene expression in relation to the secretory genes INS [I] GCG [J], SST [K], PPY [L], and IAPP [M]. Spearman’s ρ (r) and the P-value of the gene based on the empirical correlation distribution is reported. INS = Insulin, GCG = glucagon, SST = somatostatin, PPY = pancreatic polypeptide, IAPP = islet amyloid polypeptide.

**Supplementary figure 6:**
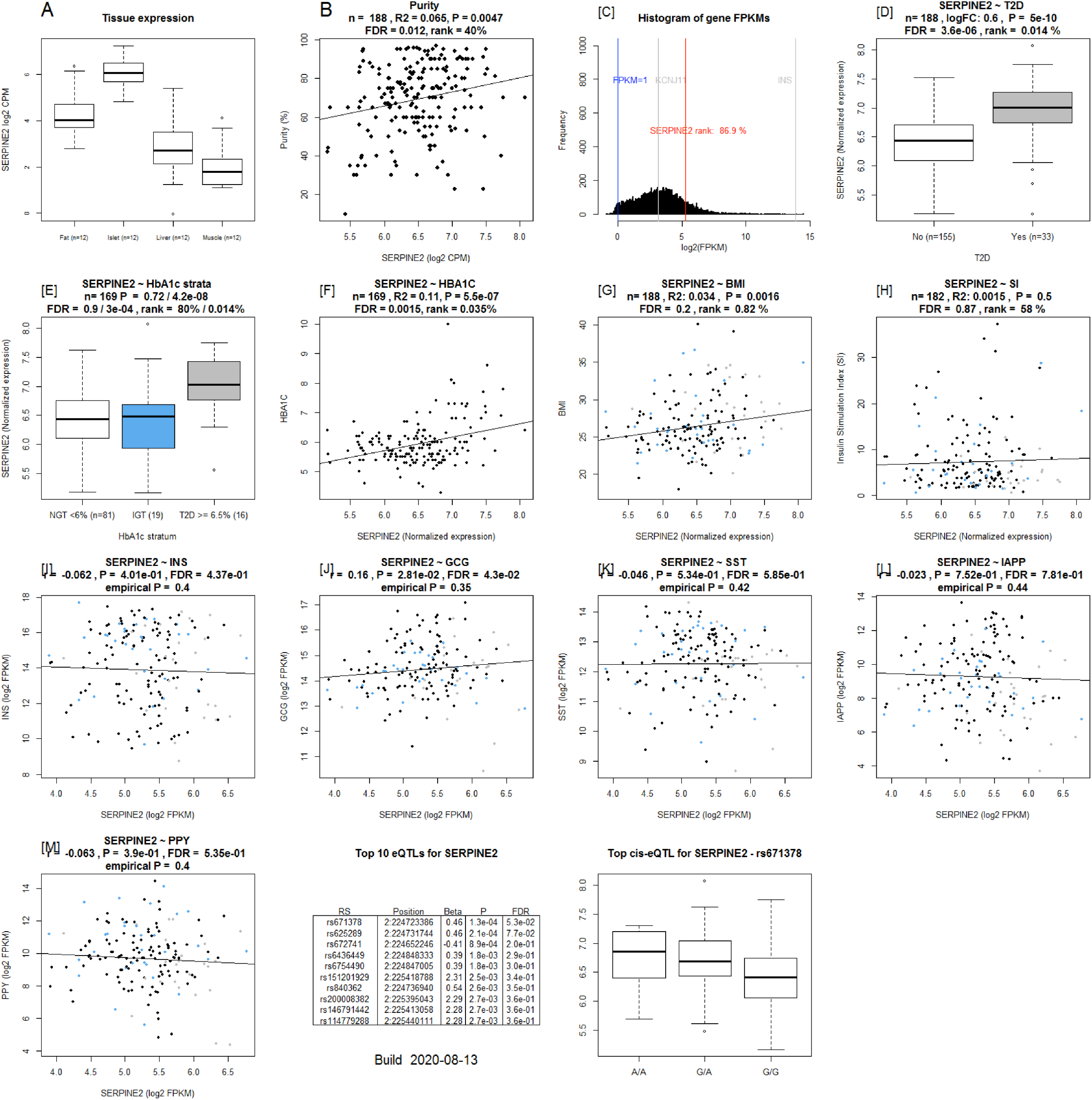
Islet Gene View graph of *SERPINE2*. shows highest expression in islets compared to in fat, liver, and muscle in the same pool of 12 individuals [A], *SERPINE2* expression is positively correlated with purity, (endocrine component) [B] and is ranked 86.9% of genes expressed in the islets [C]. *SERPINE2* expression is significantly upregulated in islets from T2D donors compared to controls [D], as well as compared to NGT donors [E] and is positively correlated with Hb1c levels [F]. *SERPINE2* expression in nominally positively correlated with BMI [G]. Test statistics are reported, namely: coefficient of determination (R2), nominal P-value, and percentage rank among all genes as calculated based on sorted P-values. [I-M]: Gene expression in relation to the secretory genes INS [I] GCG [J], SST [K], PPY [L], and IAPP [M]. Spearman’s ρ (r) and the P-value of the gene based on the empirical correlation distribution is reported. INS = Insulin, GCG = glucagon, SST = somatostatin, PPY = pancreatic polypeptide, IAPP = islet amyloid polypeptide.

**Supplementary figure 7:**
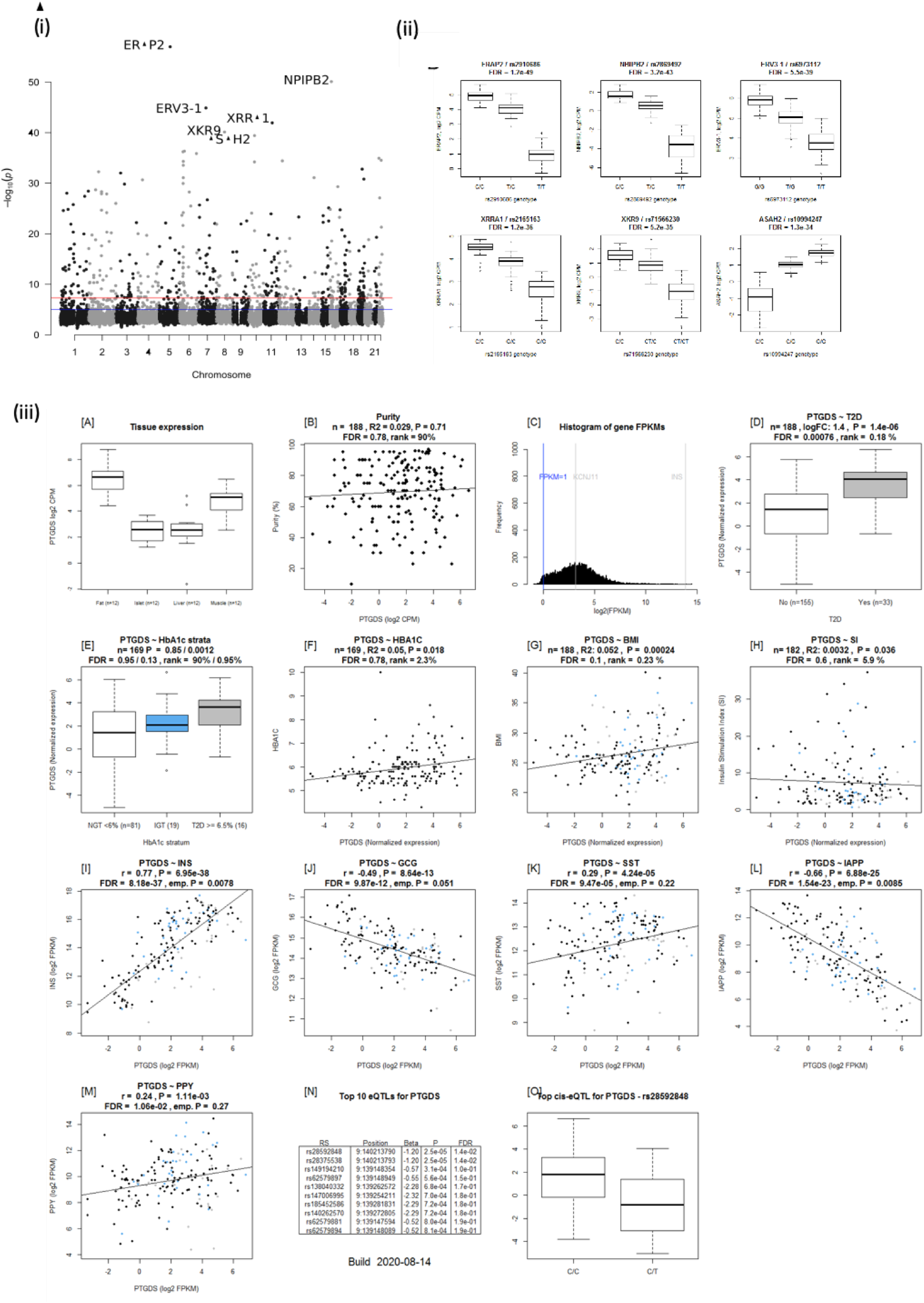
(i) Manhattan plot of eQTLs with the lowest P-value per gene. The top 6 eQTLs are marked with their target gene. (ii) Boxplots of the top 6 most significant gene-lead eQTLs. (iii) Islet geneview plot of PTDGS gene. PTGDS is expressed in fat, liver, islet and muscle, and is in the top 65.6% of all genes expressed. PTGDS expression is upregulated in islets from T2D donors, positively correlated with BMI and INS expression whereas negatively with GCG and IAPP. 2 Significant eQTLS for PTGDS are reported. The top eQTL is rs28592848, wherein T allele carriers have lower expression than C.

**Supplementary figure 8:**
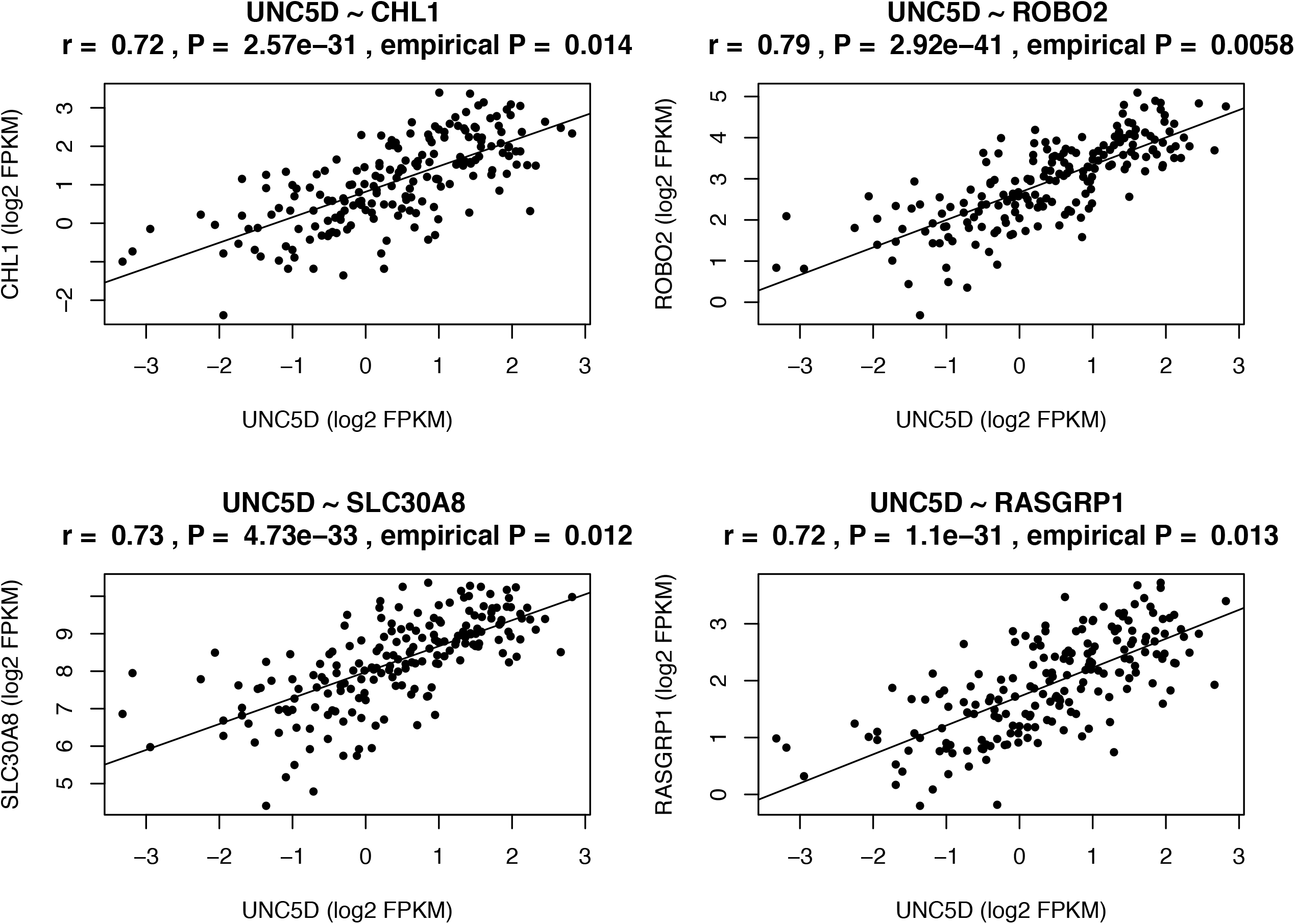
UNC5D and insulin secretion / T2D loci. UNC5D expression was significantly correlated with that of CHL1, RASGRP1, ROBO2 and SLC30A8.

**Supplementary figure 9:**
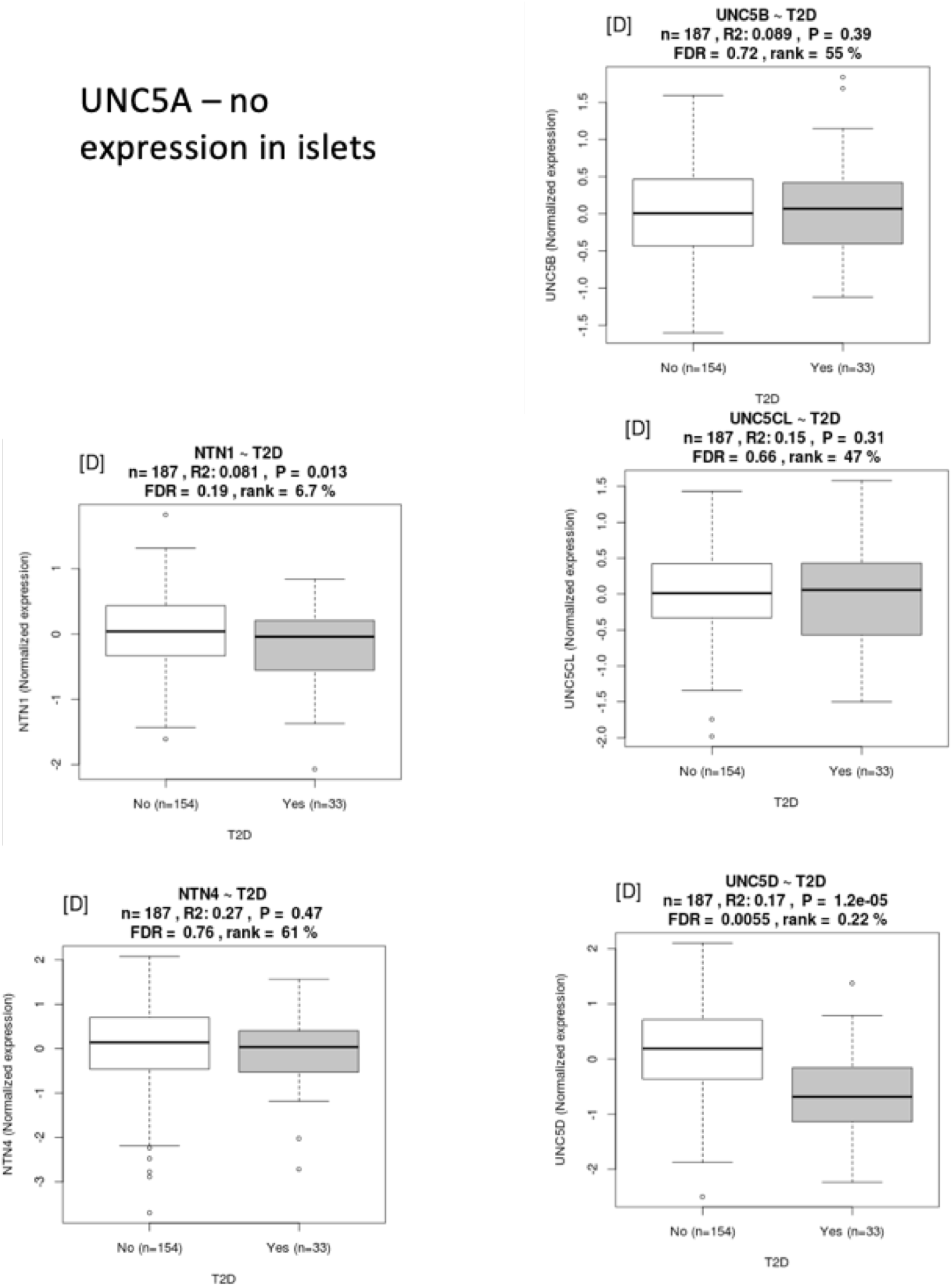
Expression of the UNC5 family and Nectrins in islets. (A) UNC5A is not expressed (B) UNC5B (C) UNC5C is shows nominal signals of downregulation in T2D donor islets (D) UNC5L (E) NTN4 and (F) UNC5D.

